# Geometric reshaping of task-relevant representations in the primary visual cortex supports perceptual decisions

**DOI:** 10.64898/2026.08.29.747961

**Authors:** Julien Corbo, Leyla Roksan Caglar, O. Batuhan Erkat, Pierre-Olivier Polack

**Affiliations:** Center for Molecular and Behavioral Neuroscience, Rutgers University–Newark, Newark, NJ, USA; Windreich Department of AI and Human Health, Icahn School of Medicine at Mount Sinai, New York, NY, USA; SUNY College of Optometry, Department of Biological and Vision Sciences, New York, NY, USA

## Abstract

Discriminating between two stimuli requires that their neural representations become separable by downstream readouts. This can be achieved geometrically, by disentangling the manifolds that population responses form in neural state space. Such reorganization has been observed in associative and motor areas, but never at the earliest stage of cortical processing. While task learning is known to impact neuronal representations in the primary visual cortex (V1), it is unknown if the population geometry is also reshaped to support perceptual decisions. We imaged V1 populations in mice trained on a Go/NoGo orientation discrimination task of increasing difficulty, and in naive mice passively viewing the same stimuli. Training reshaped the representational geometry so that the population responses were better linearly separable. A static compression made the Go and NoGo manifolds more compact and lower-dimensional from the earliest response, while a dynamic separation drove them further apart through the trial. Together, those transformations increased manifold capacity and readout accuracy. This reorganization made the Go-NoGo relationship in the neural state space more stable in Trained animals than in Naive ones. Within this learned geometry, the position of individual trials along the Go–NoGo axis predicted the animals’ decision probabilities. Learning therefore promotes a disentangled and stable representational geometry that feeds the decision process.

## Introduction

A mouse learning to tell two gratings apart, or a radiologist learning to spot a tumour on a scan, faces the same underlying problem, namely that the two objects must come to evoke reliably different responses. What has to change in the brain for this to happen and where? Learning reshapes neural representations to meet the computational demands imposed by the task at hand (1). In that case, telling two stimuli apart ultimately requires that their population responses be distinguishable by the downstream circuits reading them out. This challenge has a natural geometric description. The responses form, for each stimulus, a cloud of points in the neural state space spanning trial-to-trial variability, often termed an object manifold (2, 3). Discrimination has been proposed to be solved by pulling the two manifolds apart into linearly separable regions of the state space, such that even a simple readout, i.e. a weighted sum of the population as a down-stream neuron could implement, suffices to tell them apart (4, 5). How readily such a linear readout can separate object manifolds depends on the geometric properties of the manifolds, such as their spread (radius), dimensionality, alignment and distance in neural state space (2, 3, 6). Across systems, geometric reshaping appears to be a general mechanism by which the brain increases the linear separability of task-relevant representations, operating over multiple timescales and levels of the hierarchy. Linear classification capacity increases along the processing hierarchy - from V1 to V2, V4 to IT in the primate ventral stream, and along human ventral temporal cortex (7, 8), as well as from V1 to LM/AL/LI in the mouse (3) - mirroring the layer-by-layer gains seen in trained deep networks (9). Learning itself reshapes manifold geometry in mouse posterior parietal cortex, IT and hippocampus (8, 10), and perceptual learning shrinks and separates stimulus manifolds in human motion-selective areas (V3A and hMT+; (11). Comparable disentangling even unfolds within single trials, as motor and premotor representations progressively separate during movement planning and execution (8). Together, these findings establish geometric reshaping as a general mechanism by which the brain optimizes the linear separability of task-relevant representations, operating over multiple timescales and at multiple levels of the hierarchy.

Yet, these demonstrations have been concentrated in down-stream or higher-order areas. The primary visual cortex, the earliest stage of cortical visual processing, has not been examined from this perspective in the context of learning. Nevertheless, V1 is a natural place to look for such reorganization: rather than a passive feedforward relay, it is dominated by intracortical recurrent and cortico-cortical feedback connections, and its responses are dynamically shaped by context, expectation and task (12–15). Specifically, we know that training on a visual discrimination task substantially modifies V1 population responses: the number or responsivity of neurons tuned to task-relevant orientations increases (16–19), activity becomes sparser (20, 21), and representations of task stimuli become more distinct at the population level (17, 20–22). Whether these feature-encoding-level changes reflect a geometric reorganization of V1 population responses that supports manifold disentanglement remains unknown. Given that training already sharpens and separates V1 representations at the feature level, we hypothesized that it also reshapes their population geometry, disentangling the task-stimulus manifolds to improve their linear separability. To test this, we used two-photon calcium imaging in the layer 2/3 of V1 in mice trained on a Go/NoGo orientation discrimination task and in Naive mice passively viewing the same stimuli, across a range of Go/NoGo angular separations (90° to 15°). We characterized the geometry of Go and NoGo population response manifolds using geometric and dimensionality measures, together with direct measures of linear separability: manifold capacity and linear SVMs. We found that learning reshaped the geometric properties of the population response to the task cues, with both static and dynamic effects. Trained animals had more compact object manifolds during the entire stimulus presentation. Concurrently, the separability of their Go and NoGo representations was dynamically improved throughout the trial, contrasting with that of Naive animals that collapsed after the early response. The geometric properties were more stable across Go/NoGo angles in Trained animals, while they tracked the stimulus features in Naive animals. Hence, training constrained the geometry of the responses so the spatial relationship between the task-relevant representations was stabilized in the neural state space. Critically, the position of individual trials relative to that learned Go/NoGo geometry predicted licking probability, linking the geometric reorganization to the downstream readout that supports behavior.

## Results

We used calcium imaging data recorded in the layer 2/3 of mice trained for a Go/NoGo orientation discrimination task (n=10 *×* 6 sessions, 250 *±* 12 neurons per session), as well as Naive mice passively viewing the same stimuli (n=7 *×* 6 sessions, 475 *±* 24 neurons per session) (22). The task consisted in the randomized presentation of Go and NoGo oriented drifting gratings, and the water-restricted animals had to lick during the presentation of the Go cue (45°) to get a water reward, and refrain from licking during NoGo (135°) trials (**fig. 1a**). Trained mice underwent daily training until they reached expertise and then went through six consecutive days of recording, during which the NoGo cue was discretely made progressively closer to the Go orientation. The Go/NoGo angle went from 90° on day 1 to 15° on day 6 (**fig. 1b**). Thus, the task increased in difficulty, from the easiest possible angle (90°) to the limit of the mice’s discrimination ability (23). Animals learned the task by initially licking at detection of any stimulus, then progressively learned to withhold licking for the NoGo stimulus, demonstrating discrimination (**fig. 1c**). Their performance gradually decreased in accuracy as the Go/NoGo angle reduced. This decrease was carried by an increase in false alarms (FA), indicative of the NoGo being more and more confused as being the Go stimulus (**fig. 1d**). Neuronal activity was deconvolved (24) into Action Potential-related events (APrEs), baseline z-scored and binned in 3 frames (194ms) windows (**fig. 1e**).

**Figure 1.**
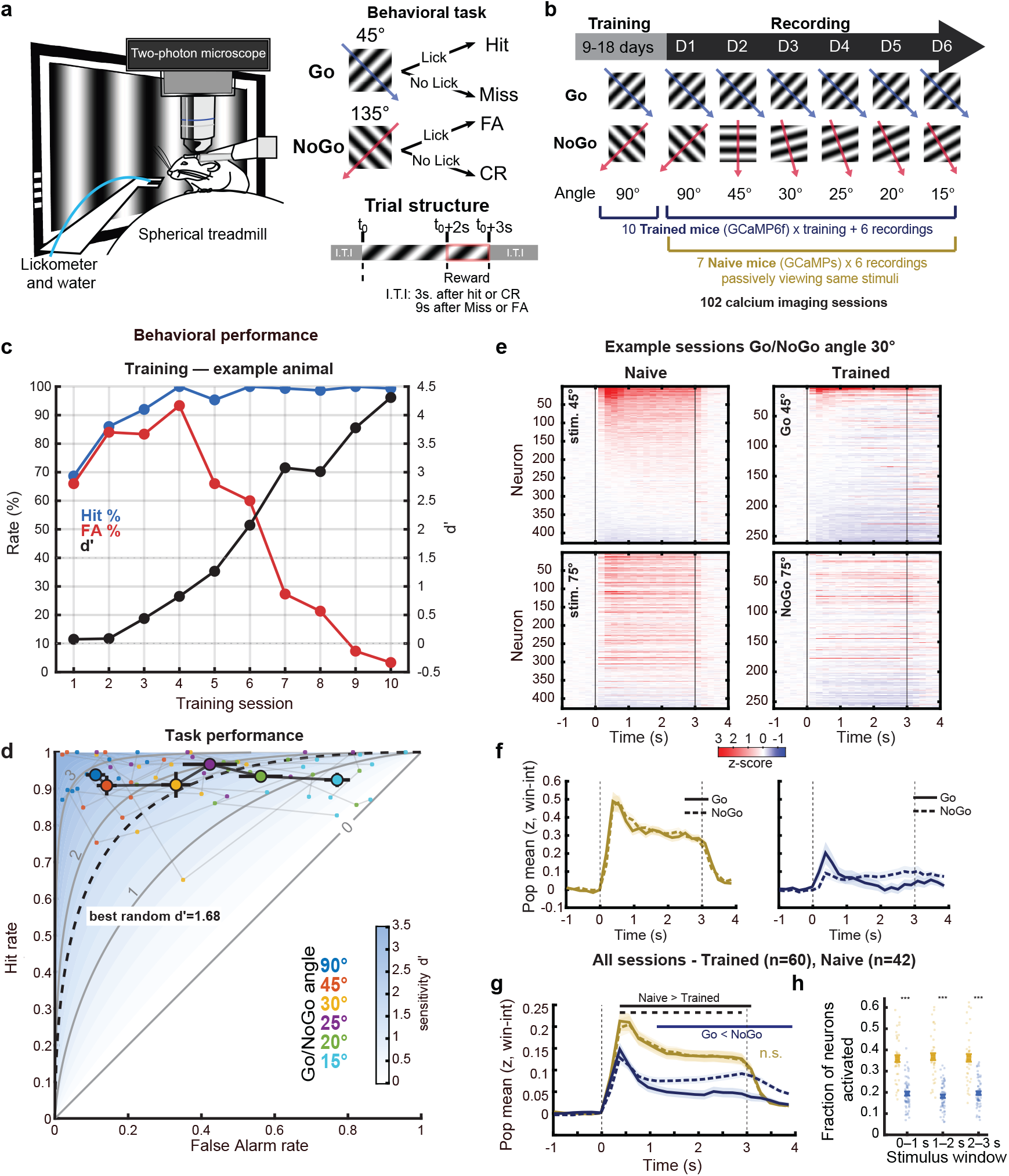
Calcium imaging recording of V1 populations in mice performing an orientation Go/NoGo task or passively viewing the same stimuli. **a**. Left panel: Schematic of the experimental setup. Right panel: Go/NoGo orientation task structure. **b**. Experimental design of the training and recording sessions. **c**. Training time course for an example mouse. Hit rate (blue), False Alarm rate (red), discrimination index D’ (black). The mouse first learned to lick for any stimulus detection (until day 4), then progressively learned to discriminate between the two cues (decrease in FA rate). **d**. Summary of the behavioral performance of the Trained cohort. Data is presented as Hit vs. FA rates, defining the D’ space. Individual sessions are indicated by small dots, connected within animals. The color indicates the Go/NoGo angle. Large dots indicate across-animal averages, with bars indicating s.e.m.. Dashed line: discrimination threshold (best random D’). **e**. Average peri-stimulus neuronal activity evoked by the two task cues in a Naive and a Trained animal. Color indicates z-score values. Every line corresponds to a neuron. **f**. Average population activity (z-score) for the example shown in e (Naive in gold, Trained in blue). Solid lines: activity evoked by the Go stimulus, dashed lines: by the NoGo stimulus. Shaded areas indicate the s.e.m. **g**. Average of all sessions pooled together (n=60 Trained, n=42 Naive), same conventions as in f. Horizontal lines indicate the time bins at which the activity was different between two time courses (permutation test with cluster correction p<0.05) **h**. Fraction of responsive neurons in 1s time windows during stimulus presentation, in Trained (blue) and Naive (gold) animals. Stars indicate FDR corrected wilcoxon ranked test p<0.001

Trained animals exhibited lower levels of population activity than Naive (**fig. 1f**; cluster-based permutation test, Naive vs. Trained, Go: cluster p < 10^−4^, significant from the second time bin post-stimulus [0.31 s] to stimulus end, sustained (1– 3 s) Hedges g = 1.05; NoGo: cluster p = 3 *×* 10^−4^, from 0.31 s, g = 0.74), showing sparser evoked responses to the Go and NoGo stimuli (**fig. 1g**; fraction of responsive neurons over the stimulus presentation: median and IQR Trained = 0.25 [0.20–0.32], Naive = 0.45 [0.39–0.53]; Wilcoxon rank test p = 3 *×* 10^−12^, g = 1.9). Moreover, the time course of activity evoked by the two stimuli was similar in Naive animals (Go vs. NoGo, n.s.), while the NoGo evoked, on average, more activity in the late response of trained animals (**fig. 1f**, paired permutation cluster p < 10^−4^ starting at 890 ms poststim, g_1-3s_ = 0.45). To determine whether this sparsening impacted the geometry such that it generated more compact and separable object manifolds, we first visualized the population responses in a lower dimension projection.

### Visualization of V1’s task representational geometry

To visualize the activity in a low-dimensional neural state space, we used PaCMAP, a nonlinear dimensionality reduction method designed to optimally maintain spatial relationships across trials within and between stimulus categories (25). We applied PaCMAP to individual recording sessions and visualized the projected point clouds in the embedding space at different time points, as well at their centroid trajectories. We found that the point clouds for the Go and NoGo stimuli started at a similar baseline location in space and quickly diverged away from their origin and from each other, before remaining at a stable location during stimulus presentation (see example **fig. 2a-b**). After the stimulus ended, the Trained point clouds stayed in their stimulus - on location for longer while the Naive point clouds almost immediately returned to their origin. Moreover, the Trained point clouds were visually more compact after stimulus onset (**fig. 2a**), and the angle between the Go and NoGo trajectories was wider (**fig. 2b**). To see if those observations generalized across animals and task cue angles, we aligned the individual point clouds using Procrustes transformations (rotation and translation only), first for all sessions/animals within a recording day (same Go/NoGo pair) and then across all days, allowing to compare all sessions across all Go/NoGo angles (**fig. 2c & S1**). The angle of the early Go/NoGo trajectory divergence was consistently wider in Trained animals (**fig. 2d**; Kruskal–Wallis, effect of group p = 1.7 *×* 10^−5^) and decreased with the Go/NoGo angle in both groups (Kruskal–Wallis, effect of day: Naive p = 2.5 *×* 10^−4^, Trained p = 2.0 *×* 10^−7^). As the separation narrowed from 90° to 30° over the first three days, the median angle fell from 79.4° (IQR [56.3, 92.6]) to 28.9° ([16.0, 68.5]) in Naive animals but stayed near-orthogonal in Trained animals (80.0° [63.1, 138.1] to 89.5° [63.9, 98.1]). Additionally, the embeddings revealed that trained animals had a more compact and stereotyped Go/NoGo point clouds across animals and Go/NoGo pairs (**fig. 2c-e**). This maintained spatial relationship of the Go and NoGo representations in Trained animals seemed to lead to less tangled object-manifolds, conserving a better separation at shallower Go/NoGo angles while their overlap grew inversely with the Go/NoGo angle in Naive animals. We made sure that the improved separation of Go and NoGo responses were not specific to the preprocessing pipeline (**fig. S2**) nor to PaCMAP, by reproducing those results using a large panel of dimensionality reduction approaches (UMATO, cMDS, nMDS, ISOMAP, **fig. S3**). Therefore, the visualization of the responses’ geometry suggests that training stabilized the task stimuli’s representations from trial to trial and orthogonalized their trajectories in the V1 neural state space, and we resolved to confirm those modifications by quantifying them in the native space.

**Figure 2.**
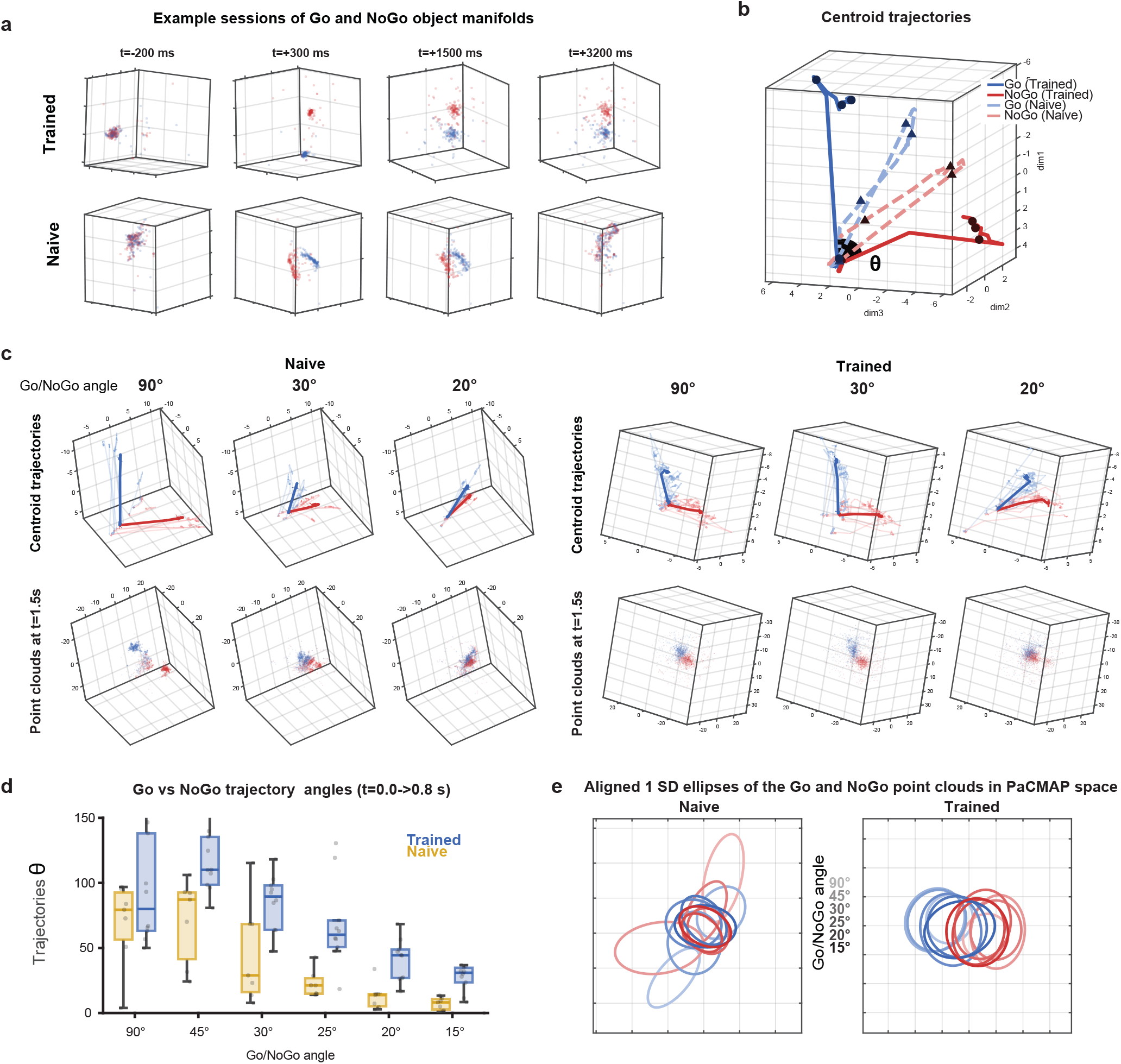
Visualization of the object manifolds with low-dimensional embeddings. **a**. PaCMAP projection of the Go (blue) and NoGo (red) single trials point clouds in the neural state space for an example Trained (top) and Naive (bottom) animal recorded at a Go/NoGo angle of 30°. Different snap shots are visualized, from left to right at t=−200ms, +300ms, +1500ms and +3200ms relative to stimulus onset. **b**. Trajectories of the point clouds’ centroids from −500ms to 3500ms relative to stimulus onset, for Go and NoGo of the two sessions from a. Darker plain lines indicate the trajectory of the Trained animal’s responses, dashed lighter lines that of the Naive animal. Black markers show the time of the snapshots in a. The angle between the Go and NoGo trajectories is indicated as theta. **c**. Procrustes-aligned trajectories and point clouds for all Naive (n-=7) and Trained (n=10) sessions at Go/NoGo angle of 90°, 30°and 20°respectively from left to right. **d**. Trajectory angles summary, computed as the angle between the Go and NoGo point cloud centroid displacement from t=0 to t =800ms in every animal and Go/NoGo angle. **e**. Ellipses fit to 1 standard deviation around the centroid of the pooled Go (blue) and NoGo (red) point clouds in the 2D plane maximizing Go/NoGo separation for Naive (left) and Trained (right) animals. Darker shades indicate smaller Go/NoGo angles.

### Training reshapes the geometric properties of task stimulus manifolds

The PaCMAP visualizations revealed qualitative differences in the compactness and spatial organization of Go and NoGo representations between Naive and Trained animals (Fig. **2a–d**). To quantify these differences, we characterized the geometry of the population response manifolds directly in the native neural state space, without relying on dimensionality reduction methods whose algorithmic choices can distort the very geometric properties we seek to measure. We characterized three geometric properties of the manifolds: their centroid distance (the separation between the mean Go and NoGo population responses), their radius (the mean distance of individual trials to the class centroid, capturing the spread of the response cloud), and their dimensionality (the number of independent axes each response cloud spans). Together, these properties shape how readily a linear readout can separate the two stimulus classes. Geometric frameworks predict that disentangling is achieved via an increase in across-class centroid distance, a decrease in within-class radius, and a reduction in dimensionality (7), which we individually quantify below.

The time course of the Go and NoGo object manifolds’ centroid distance revealed a pronounced difference in the late response dynamics between groups (**fig. 3a–d & S4)**. Both Naive and trained animals exhibited a large early increase in Go/NoGo centroid distance at stimulus onset. Expectedly, that centroid distance peak decreased with the Go/NoGo angle (**fig. 3a**, effect of day on peak, Kruskal–Wallis: pTrained = 5.0 *×* 10^−4^, pNaive = 0.022). After the early response, however, the trajectories diverged: centroid distances decayed back toward baseline in Naive animals, while they remained elevated in trained animals (**fig. 3b**). This divergence followed the same dynamic regardless of the Go/NoGo angle (**fig. 3c**). Pooling all sessions together (**fig. 3d**), we found the centroid distance between Go and NoGo representations to be higher in trained than Naive animals over the late response window (1–3 s; 0.52 *±* 0.03 vs 0.41 *±* 0.03 on average; effect size Hedges g = 0.56). The divergence between trained and Naive was significant starting 1.28s post-stimulus onset (cluster-based permutation test, significant from t ≈1.28 s post-onset and sustained through the end of the trial, p = 0.004, g>0.2). This maintained distance between centroids in trained animals, while it decreases in Naive animals, suggests that training promoted a better response separation in the neural state space, dynamically throughout stimulus presentation.

**Figure 3.**
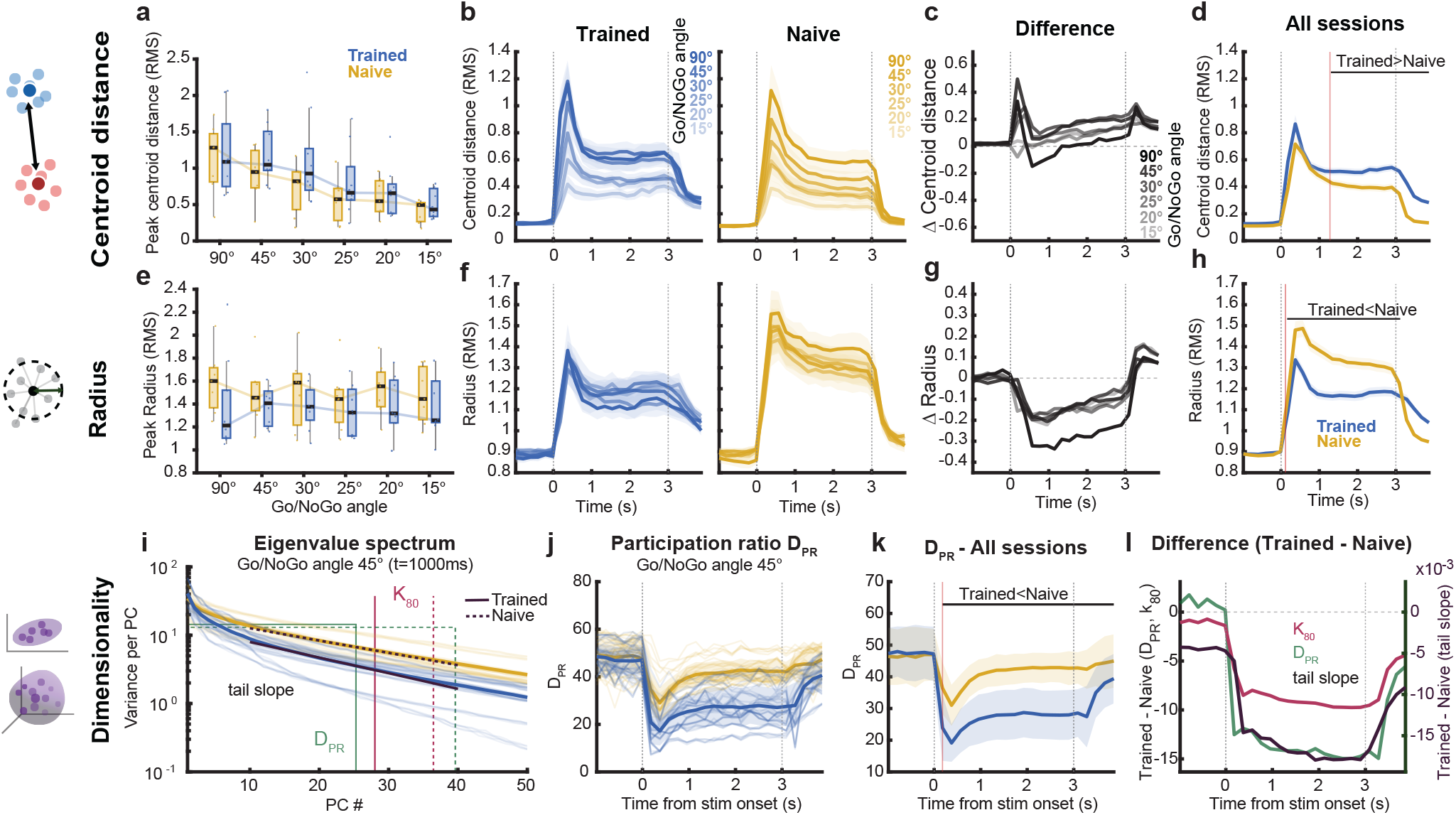
Population responses in Trained animals are more distant and compact in the neural state space. **a**. Peak centroid distance (RMS-normalized euclidean distances) summary in the native space (no dimensionality reduction) for Trained (blue) and Naive (gold) animals, across Go/NoGo angles. **b**. Time course of the average centroid distance between Go and NoGo for every Go/NoGo angle (darker shades indicate larger Go/NoGo angles), in Trained (left) and Naive (right) animals. Shaded areas indicate s.e.m. **c**. Difference in centroid distance between Trained and Naive animals for every Go/NoGo angle (darker shades indicate larger angles) **d**. Pooled centroid distance average over all sessions for Trained (blue) and Naive (gold) animals. Horizontal line indicates significant difference between groups, vertical line indicates the earliest significant difference. **e. through h**. Same as a-d but for the point cloud radii computed as the mean distance of every trial to its within-class centroid. **i**. Eigenvalue spectrum (variance explained per principal component) at t=1000s post stimulus onset for all Trained (blue) and Naive (gold) sessions on an example Go/NoGo angle (45°). Thin lines show individual sessions, thick lines show group averages. The three dimensionality measures are overlaid for each group (solid lines: Trained; dashed lines: Naive). Green: the equivalent flat spectrum — equal-variance components with the same total variance and the same participation ratio as the data — whose width is DPR. Red: K80, the component at which 80% of the cumulative variance is reached. Dark purple: the log-eigenvalue tail slope, fit over components 10–40. All overlays show group means at this snapshot. **j**. Participation ratio (DPR) time course for the same example Go/NoGo angle. Thin lines indicate values for single sessions, thick lines the within-group average, with the s.e.m. shown as shaded areas. Horizontal line indicates significant difference between groups, vertical line indicates the earliest significant difference. **k**. Same as j. but pooling all sessions from all Go/NoGo angles together, showing their grand average. **l**. Difference between Trained and Naive pooled averages (negative values indicate a lower Trained dimensionality) for all three dimensionality measures : DPR (blue), k80 (red) and tail slope (green).

Manifold radius showed a complementary, angle-invariant effect throughout stimulus presentation (**fig. 3e–h & S4**). We first found that the manifold radius did not vary with the Go/NoGo angle, indicating the absence of a stimulus proximity-induced modulation within sessions (**fig. 3e**, effect of day on peak radius, Kruskal–Wallis: pNaive = 0.88, pTrained = 0.92). Trained animals nonetheless had consistently smaller radii than Naive animals at every Go/NoGo angle (**fig. 3f–g**). Pooling all sessions (**fig. 3h**), the trained radius was lower from t ≈0.12 s after stimulus onset and sustained through the stimulus presentation (cluster-based permutation, p = 1 *×* 10^−4^, g > 0.2). Over the sustained window, mean radius was lower in trained (1.17 *±* 0.02) than Naive animals (1.32 *±* 0.03;1–3 s mean; rank-sum p = 1.2 *×* 10^−4^, g = −0.86), indicating more compact, less variable trial-to-trial stimulus representations.

We next asked whether training also reduced the dimensionality of these manifolds. We characterized their spectral dimensionality - the effective number of independent axes over which the population responses vary, read from the eigen-spectrum of their covariance - at two levels. First, we quantified the overall dimensionality of the population response, measured as the participation ratio (DPR) of the covariance spectrum pooled across all trials and stimulus classes. Overall spectral dimensionality was significantly lower in trained than Naive animals (1–3 s window; Trained 39.5 *±* 1.6 vs Naive 62.7 *±* 1.2, mean SEM; Wilcoxon rank-sum p = 1.4 10^−14^, g = −2.14), indicating that learning reduces the overall dimensionality of V1 population responses. Because this measure confounds within-class variability with between-class separation, and because trained animals show larger centroid distances (see **fig. 3a-d**) - which inflates the between-class component and pushes overall dimensionality upward - the observed reduction should be driven by a compression of the individual stimulus clouds that is strong enough to overcome the between-class inflation. To isolate this within-class compression, we turned to the second level: the dimensionality of the individual stimulus clouds, measured as the participation ratio of the within-class covariance during the time course of the trial (Fig. **3i–j**). Again, the participation ratio of the within-class covariance was significantly lower in trained animals during the stimulus presentation (cluster-based permutation test, cluster p = 10^−4^, significant from the first post-stimulus window [0.18 s] through the end of the trial), establishing that the dimensionality reduction reflects the compactness of individual stimulus representations rather than their separation. Over the sustained window (1–3 s), participation ratio was 42.5 *±* 0.7 (mean SEM) in Naive animals and 28.0 *±* 1.0 in trained animals (rank-sum p = 6.4 *×* 10^−15^, g = −2.22). This difference was consistent across all six recording days (**fig. S5**). This compression was not confined to the dominant principal components. Both k80 (the number of components required to explain 80% of within-class variance) and the log-eigenvalue tail slope were also significantly lower in trained animals across the full stimulus epoch and across all six recording days (k80: cluster p = 10^−4^ from 0.18 s; 1–3 s: 28.7 *±* 0.9 vs 38.1*×* 0.5, rank-sum p = 1.8×10^−11^, g = −1.59; tail slope: cluster p = 10^−4^ from 0.18 s; 1–3 s: 0.057 0.004 vs −0.039 *±* 0.001, rank-sum p = 3.3 *×* 10^−11^, g = −0.78; **fig. 3k**), indicating that training reshapes the entire eigenspectrum from the dominant components through the residual variance tail rather than concentrating variance only in the first few components. Notably, dimensionality dropped significantly from the pre-stimulus baseline to the stimulus epoch within both groups (paired signed-rank test, all p ≤ 2.2 *×* 10^−3^), but the stimulus-evoked compression was markedly greater in trained animals, with participation ratio falling from an equivalent pre-stimulus baseline from an equivalent pre-stimulus baseline (Naive: 46.6 *±* 1.3, Trained: 47.5 *±* 1.1; all baseline group differences p > 0.62) to substantially lower stimulus-epoch values (Naive: 42.5, Trained: 28.0 over 1–3 s). The convergence of these within-class estimators - participation ratio, k80, and tail slope points to a consistent conclusion that perceptual learning in V1 compresses the dimensionality of stimulus representations across the entire eigenspectrum, from the dominant principal components through the residual variance tail, and does so specifically during the stimulus epoch rather than as a persistent resting-state difference.

Altogether, we observed static and dynamic changes to the geometry of the trial responses in Trained animals. The reduction of the radius and dimensionality started at the earliest response (0.12s) and was static throughout the entire evoked response, while the centroid distance was improved after the early response (1.28s), ramping up as the distance in Naive animals went down. They contribute to a coherent reshaping of the responses. More compact, further apart object manifolds are predicted to be more separable, so we next wondered whether and how these different changes would synergize to improve the linear separability of the Go and NoGo responses.

### The geometric changes in Trained animals enhance linear separability of task stimulus representations

The geometric changes described above - increased centroid distance, reduced radius, and lower dimensionality - are each predicted to improve linear separability, but the individual measures do not tell us whether and how they combine to make the two classes more separable overall. To assess this directly, we computed the manifold capacity of the Go and NoGo population responses (**fig. 4a-f & S6**). Manifold capacity is a framework that captures the efficiency of the geometry (radius, centroid distance, dimensionality and alignment) of the different classes’ representations with respect to their linear separability. It essentially asks how many objects an average neuron can linearly separate if the objects’ representational manifolds are shaped as the ones in the data. With higher capacity indicating more separable representations, this measure therefore allows to directly relate the geometric reshaping to the improved separability. Here we estimated capacity directly from the geometry of the two manifolds rather than from summary statistics of their radius, dimensionality, and position (see Methods), and computed it per session from the full population using all available trials.

**Figure 4.**
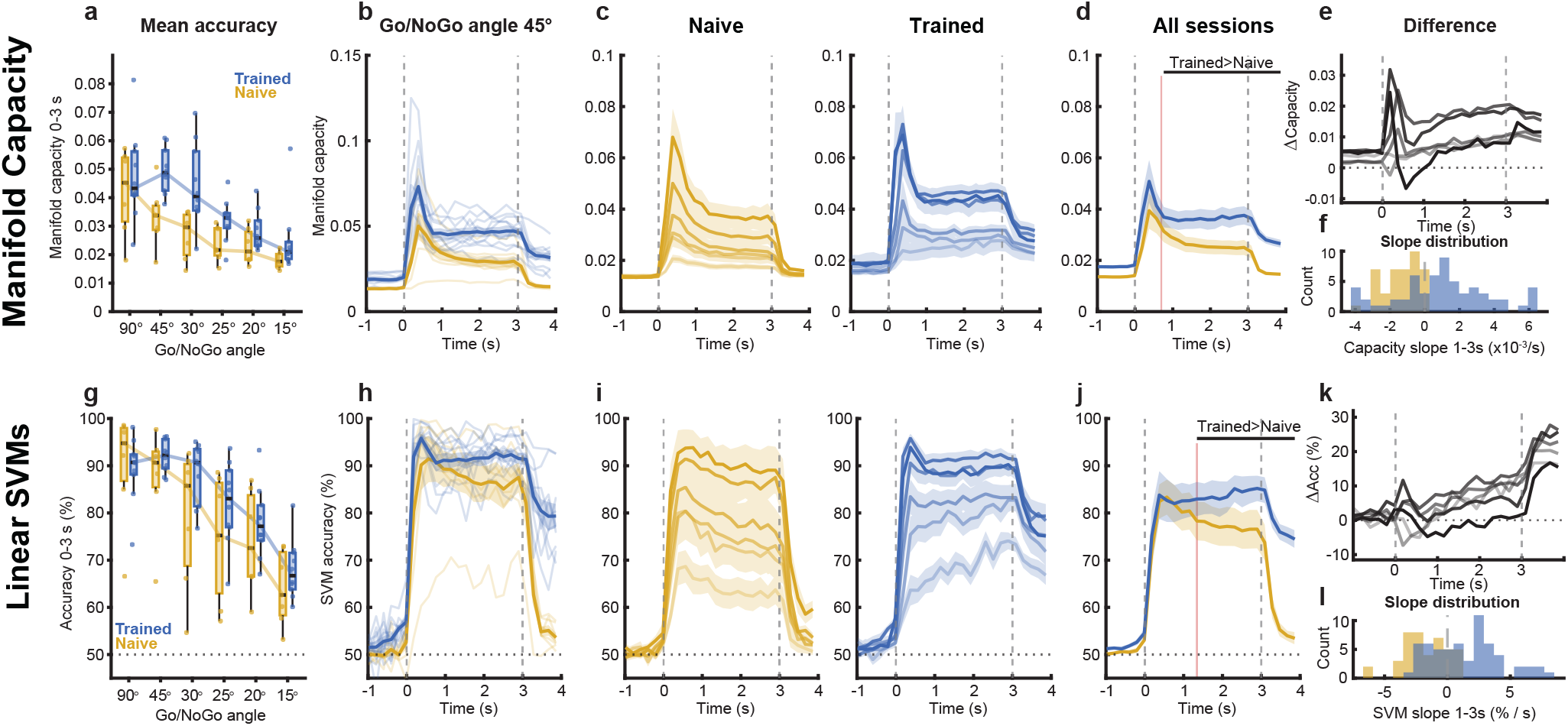
Training improved the linear separability of the responses to the task stimuli. **a**. Manifold capacity averaged over the whole stimulus presentation across Go/NoGo angles in Trained (blue) and Naive (gold) animals. **b**. Manifold capacity time course for an example Go/NoGo angle (45°) using Trained (blue) and Naive (gold) population responses. Single sessions are shown as thin lines and group averages as thick lines, with the shaded area indicating s.e.m. **c**. Time course of the average manifold capacity for every Go/NoGo angle (darker shades indicate larger Go/NoGo angles), in Trained (left) and Naive (right) animals. Shaded areas indicate s.e.m. **d**. Pooled manifold capacity averaged over all sessions for Trained (blue) and Naive (gold) animals. Horizontal line indicates significant difference between groups (permutation test with cluster correction), vertical line indicates the earliest significant difference. **e**. Difference in manifold capacity between Trained and Naive animals for every Go/NoGo angle (darker shades indicate larger angles). **f**. Distribution of single sessions manifold capacity slopes between 1s and 3s after stimulus onset for the Trained (blue) and Naive (gold) cohorts. **g. through l**. Same as a through f but for the linear SVMs trained on Naive and Trained populations.

We found that manifold capacity was higher at wider Go/NoGo angles in both groups (**fig. 4a**, 0–3 s average capacity at Go/NoGo angle 90°: Trained = 0.047 *±* 0.016, Naïve = 0.042 *±* 0.015) and decreased as the stimuli became closer (0–3 s average capacity at Go/NoGo angle 15°: Trained = 0.025 *±* 0.012, Naive = 0.018 *±* 0.003; effect of Go/NoGo angle on capacity, Kruskal–Wallis pNaive = 0.0051, pTrained = 9.4 *×* 10^−6^). The capacity time course revealed different dynamics between the two groups (**fig. 4b–f**). After a similar early transient, Trained capacity increased (1–3 s slope, median +0.0011 per s, IQR [−0.0005, 0.0026]) while Naive capacity decreased consistently (median −0.0011 per s, IQR [−0.0020, −0.0005]; Naive vs Trained slope, rank-sum p = 2.3 *×* 10^−7^). Averaging over all sessions, Trained capacity exceeded Naive capacity from 0.70s after stimulus onset (cluster-based permutation test, p= 1.0 *×* 10^−^4; **fig. 4d**). Because manifold capacity is normalized per neuron, this advantage cannot be attributed to population size, and because it is measured directly rather than derived from our geometric measures, its agreement with them reflects convergent evidence that the geometric reorganization is sufficient to improve separability.

Manifold capacity shows that the manifolds *can* be separated given their geometry. We next asked whether a learned decoder realizes this and generalizes it to unseen trials. To test this with a simple linear readout mechanism (26) we trained linear Support Vector Machines (SVMs) to classify Go versus NoGo responses and evaluate them on held-out data (five-fold cross-validation; **fig. 4g–l & S6**). To make sessions comparable, neurons were subsampled to a common ceiling (250 or all available per session; Trained N = 211*±* 47, Naive N = 249 *±* 7) and trials balanced across classes (up to 100 per class). Accuracy was high at wide Go/NoGo angles in both groups (**fig. 4g**, 0–3 s average performance at 90°: Trained = 89.4 *±* 6.8, Naive = 90.4 *±* 11.5) and fell with stimulus proximity (0–3 s average at 15°: Trained = 68.0 *±* 6.2, Naive = 64.0 *±* 7.6; effect of Go/NoGo angle on accuracy, Kruskal– Wallis pNaive = 0.0043, pTrained = 4.1 *×* 10^−6^). As with capacity, the groups diverged over the trial time course of SVM accuracy differed between groups (**fig. 4h–l**): Trained accuracy ramped up after the initial transient (**fig. 4l**, 1–3 s slope, median +1.68 %/s, IQR [−0.84, 3.34]) while Naive accuracy declined (median 1.93 %/s, IQR [−3.01, −0.33]; Naive vs Trained slope, rank-sum p = 1.1 *×* 10^−8^), becoming steadily greater than Naive after the early response of 1.34 s post-stimulus onset (**fig. 4j**, cluster-based permutation test on all sessions pooled, p = 3.0 *×* 10^−4^). Notably, this readout advantage emerged despite the trained classifiers using, if anything, fewer neurons than the Naive ones, making the comparison conservative. A concrete linear readout thus confirms the separability advantage predicted by the manifold geometry, sustained across the trial.

### The geometric improvement is not driven by behavioral engagement in Trained animals

Having shown how training changed V1’s geometry, we next inquired about the relationship between the geometry-driven separability improvement and the behavioral performance. First, we needed to confirm that the changes to the Trained animals’ V1 geometry related to the representation of the task stimuli, and not merely to the difference in behavioural engagement between Trained and Naive animals. Performing the task indeed requires attention and motor responses, which could contribute, at least in part, to the improved geometry and linear separability. Locomotion and arousal have been shown to modulate V1 activity (27–30). Specifically, orofacial movements, which are numerous during a licking response, constitute a strong driver of V1’s neurons (31–33). Therefore, since capacity encapsulates the ensemble of the geometry-related effects we uncovered, we first compared the time course of capacity at different levels of licking activity. If licking and the associated orofacial movements were drivers of the geometric improvement, the capacity computed from trials with high licking should be greater than the capacity computed from trials with low licking. To retain enough trials to reliably compute the capacity of the Go/NoGo representations, we split the trials in tertiles computed from the area under the curve of the licking trace between 1s and 2s post-stimulus (T1-3, with highest licking in T3). Individual sessions showed large, if variable, differences in licking between the low (T1) and high (T3) tertiles, but only small accompanying differences in capacity (**fig. 5a,b**). Across the cohort, licking activity nearly doubled from the low to high tertile (**fig. 5d**; median +79%, IQR [63, 103]%, signed-rank p = 1.6 *×* 10^−11^, g = 1.80). Yet, capacity changed only marginally and, if anything, decreased (median −5.1%, IQR [−10.6, +0.5]%). The difference was statistically detectable (p = 2.7 *×* 10^−5^) but negligible in magnitude (g = −0.15), and in the direction opposite to what we would have expected from a licking confound. Consistent with this, the capacity change across sessions was weakly negatively correlated with the licking change (**fig. 5c**, r = −0.39, p = 0.0023). This relationship was not robust, as excluding the single most influential of 60 sessions rendered it non-significant (r = −0.24, p = 0.06). Thus any residual effect countered rather than supported the hypothesis that licking underpins the geometric improvement. Consistently, computing the capacity on trials split in a binary licking/no licking confirmed, when enough trials of the two classes were available, that the presence of licking, regardless of amplitude, was not the driver of the improved capacity (**fig. S7**). We performed a similar analysis on the locomotion during the same 1s-2s window post stimulus onset, and found no effect of the locomotion tertile on capacity (**fig. 5d & S8**, locomotion change median +149%, IQR [141, 162]%, p = 8.9 10^−5^; g = 0.89; capacity change median +1.4%, IQR [1.5, +5.5]%, p = 0.25, g = 0.02). The same approach applied to the pupil size (a proxy of arousal) also showed no effect on capacity when measuring either (1) the pre-trial pupil size to gauge the state of the animal going into the trial (**fig. 5d & S9**, pupil change median *+28%, IQR [22, 38]%, p = 5*.*3 × 10*^*-8*^, *g = 0*.*61*, capacity change median +2.6%, IQR [−4.2, +7.2]%, p = 0.094 (g = 0.03) or (2) the 1s-2s pupil size to gauge the trial-related modulation (**fig S10**). For both locomotion and pupil size, the asymmetry between Go and NoGo trial was also not driving capacity, since computing capacity from equalized trial pools did not change the values (**fig. S9-10**). To summarize, variations in licking, locomotion and pupil size associated with task performance lead to almost no capacity variations and could not explain the improved geometry observed in trained animals (Trained - Naive capacity median +22%, IQR [11, 38]%, rank-sum p = 1.0 *×* 10^−6^, g = 0.97). Altogether, training-related changes were about four-fold larger than the licking-related change, and more than an order of magnitude larger than the locomotion and pupil effects (**fig. 4d**).

**Figure 5.**
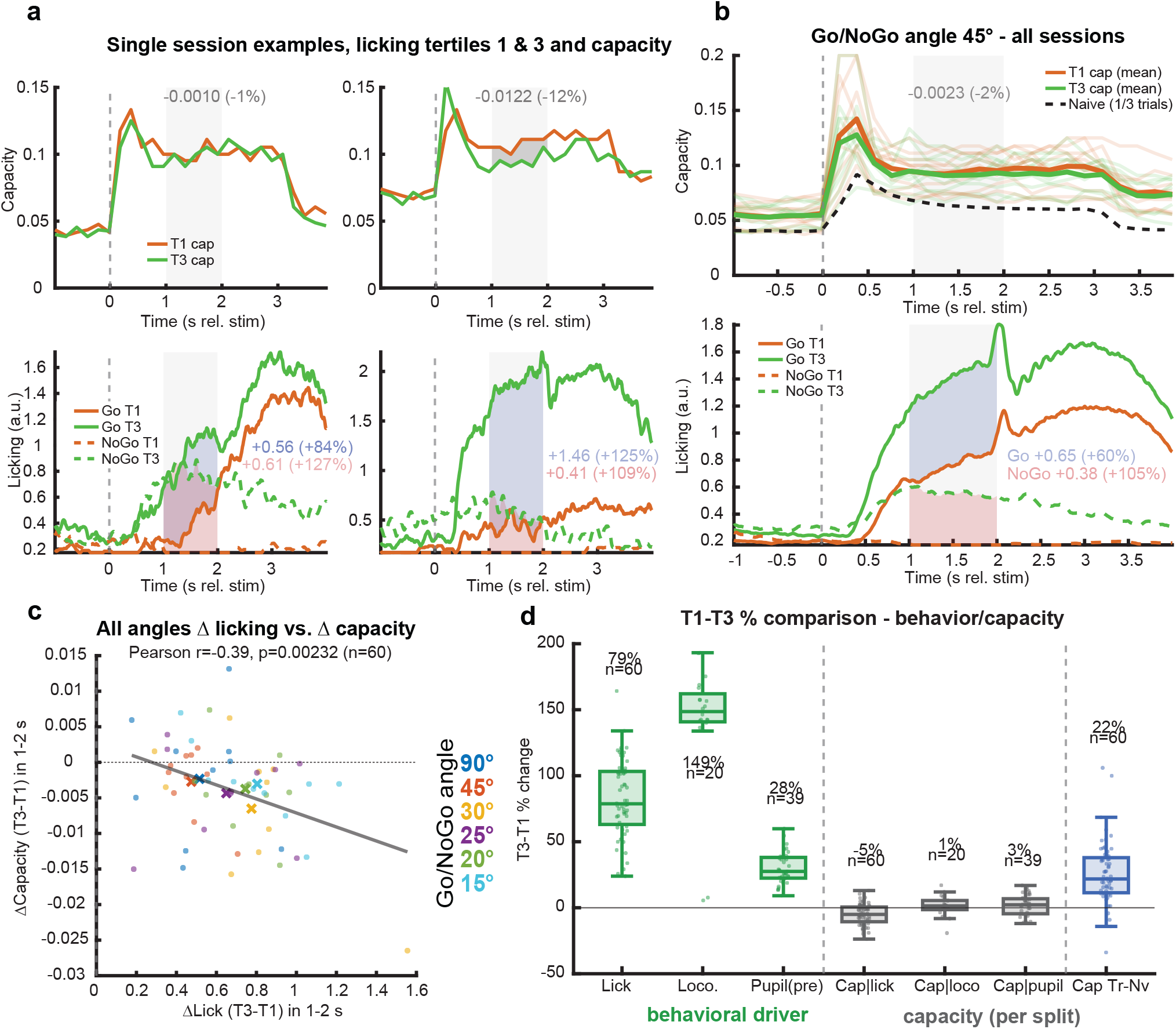
Training induced changes are not explained by behavioral engagement. **a**. Example single sessions illustrating the licking tertile control. Trials were split within each stimulus class into low (T1) and high (T3) tertiles of licking activity (integral of the lick trace over the 1–2 s window), and manifold capacity was recomputed within each tertile. Top: capacity time course for T1 (orange) and T3 (green); the T3−T1 difference averaged over the 1–2 s window is indicated (absolute and %). Bottom: licking time courses for Go (solid) and NoGo (dashed) trials in each tertile (T1 orange, T3 green). Shaded areas indicate the 1–2 s window and the tertile differences. **b**. Same analysis pooled across all sessions at a Go/NoGo angle of 45°. Top: mean capacity for the T1 (orange) and T3 (green) tertiles, with the naive level (dashed black, computed on matched 1/3-trial subsets); the T3−T1 window difference indicated. Bottom: mean Go (solid) and NoGo (dashed) licking for each tertile. Thin lines show individual sessions, thick lines the group mean; shaded areas indicate s.e.m. and the 1–2 s window. **c**. Mean T3−T1 change in capacity (1–2 s) against the mean T3−T1 change in licking (1–2 s), across all Go/NoGo angles. Each dot is a session; color indicates the Go/NoGo angle.. **d**. T3−T1 percentage change per split. Green: the behavioral variables — licking, locomotion and pre-stimulus pupil. Grey: capacity computed within each split — by licking, by locomotion and by pupil. Blue: the trained-versus-naive capacity difference. Boxes show median and interquartile range, whiskers the range, dots individual sessions.

### Changes in geometry precede perceptual discrimination behavior

We have shown that training changed the geometry of V1 responses to the task stimuli in a way that favors their linear separability, an improvement theoretically linked to an easier downstream readout (4, 6). As we showed that this improvement is not a trivial consequence of the movements and arousal of the trained animals, the reorganization must reflect changes in the task cue representations themselves. Therefore, we wondered if the improved object manifold separability could support the Trained animals’ behavioral performance by feeding the decision process. To address this, we first needed to confirm that it is mechanically possible - does the Trained geometry differ from the Naive before the decisions are finalized? As we do not have access to the true decision time of the animals, we looked for proxies in the behavioral expressions of the perceptual decisions. We observed that, if most animals tended to start licking shortly after the stimulus onset (fig. S11a-b, median time to reach half-peak licking in Go trials Got50 = 0.72s, IQR [0.60, 0.86], n = 40/60 sessions), some started licking closer to the decision window (fig. S11a-b, Got50 = 1.61s, IQR [1.45, 1.75], n = 20/60 sessions) suggesting two distinct strategies. However, in both cases, licking in Go and NoGo trials started simultaneously, then NoGo licking slowed down, separating from the Go licking activity (fig. 6a-b). We treated that separation as the earliest behavioral expression of a decision being made. The Go-NoGo licking difference onset was modulated by task difficulty, occurring later as the Go/NoGo angle shrunk (earliest at Go/NoGo angle 90°, median 0.53 s, IQR [0.49, 0.63]; latest at 15°, median 1.72s, IQR [1.28, 2.08]; Kruskal-Wallis effect of day p = 1.3×10-4). This dependency is compatible with an evidence accumulation process where the more discrimination is difficult, the later the decisions are expressed (34, 35). Because the animals tended to start licking when detecting any stimulus, then stop licking when identifying the NoGo stimulus, we estimated an upper bound of the decision times as being the last lick in correct rejection trials. In those trials, the animals definitively decided that a trial was a NoGo after starting to lick earlier. This upper bound happened later than the Go/NoGo licking separation, and was not significantly dependent on the task difficulty (**fig. 6a-b**, global median 1.22 s, IQR [0.99, 1.46], Kruskal-Wallis p = 0.07). Compared to these decision proxies, the compression of radius and dimensionality was immediately present in trained animals from the beginning of the stimulus presentation (**fig. S11e**, dimensionality onset 0.12 s, 95% CI [0.12, 0.12]; radius onset 0.12s, 95% CI [0.12, 0.31]), as well as the trajectories being more orthogonal in Trained animals (**fig. 2d**). On the other hand, the centroid distance diverged between Naive and Trained animals after the early response and kept increasing during the whole trial (**fig. S11e**, onset 1.28s, 95% CI [0.12, 2.63]). As a unifying measure of the latter ones, capacity diverged between Trained and Naïve animals around 0.70s after stimulus onset (**fig. 6b**, 95% CI [0.12, 1.28]), i.e., before and during the decision times established by the licking divergence and last CR lick proxies. These timings suggest that the static and dynamic geometric modifications occur early enough during the trials to inform the decision process.

**Figure 6.**
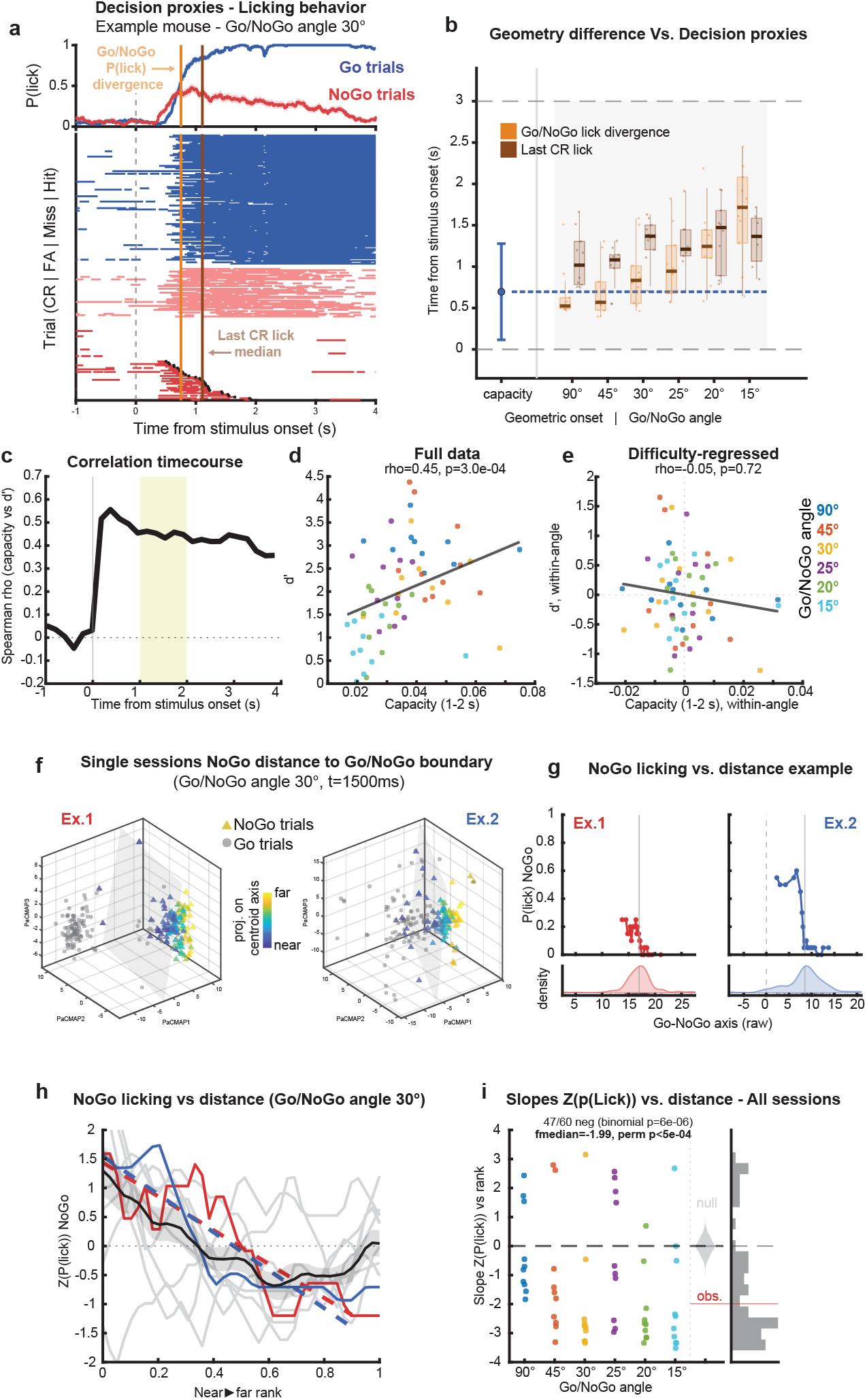
The geometry of the responses to task stimuli feed the decision process. **a**. Licking decision proxies for an example mouse at a Go/NoGo angle of 30°. Top: lick probability across trials for Go (blue) and NoGo (red) trials; shaded areas indicate s.e.m. Bottom: single-trial lick raster (Trials ordered by outcome CR (deep red) / FA (light red) / Miss / Hit (blue)); black dots mark the last lick of correct-rejection trials; dashed grey line indicates stimulus onset. The Go/NoGo P(lick) divergence (orange) and the median last correct-rejection lick (brown) are indicated as vertical lines. **b**. Geometric group divergence onset compared with the two decision proxies. Left: naive-versus-trained onset of the manifold capacity difference (dot, mean; bar, 95% boot-strap interval). Right: per-Go/NoGo angle distributions of the Go/NoGo lick divergence (orange) and the last correct-rejection lick (brown) across trained sessions; dashed blue line marks the capacity onset. Boxes show median and 25-75% quartiles, whiskers show the range, dots show individual sessions. **c**. Time course of the Spearman correlation between manifold capacity and behavioral sensitivity D^*′*^ across trained sessions; shaded band marks the 1–2 s window. **d**. D^*′*^ against manifold capacity averaged over 1–2 s, one dot per session, colored by Go/NoGo angle; line is the linear fit. **e**. Same as d after regressing out task difficulty (within-recording-day-centered capacity and d^*′*^). **f**. Singletrial NoGo position along the Go to NoGo centroid axis for two example sessions at a Go/NoGo angle of 30°, shown in the 3D PaCMAP embedding at the 1.5 s snapshot. NoGo trials (triangles) are colored by their projection onto the Go to NoGo centroid axis (blue, near the Go end; yellow, far); Go trials are in grey; the grey plane shows the Go/NoGo separating hyperplane (drawn for illustration). **g**. NoGo lick probability as a function of NoGo axis position (raw projection) for the two example sessions in f. Top: binned P(lick) on NoGo trials (sliding window of 20 trials, step 5). Bottom: density of NoGo trials along the axis; ticks mark individual trials; vertical line indicates the Go centroid. **h**. NoGo licking versus axis position aggregated across trained sessions at a Go/NoGo angle of 30°(n = 10). Each session’s binned NoGo lick probability was z-scored and plotted against the within-session near to far rank; thin grey lines show individual sessions, the black line shows the mean, and the red and blue lines the two example sessions from f–g, with dashed lines indicating their linear fits over 0-90% of the rank range. Distribution of per-session slopes of z(P(lick)) versus near to far rank, by Go/NoGo angle. The observed median (red line, “obs.”) is compared with a shuffled-null distribution (grey violin). Right: histogram of all session slopes.

### The location of activity in the space defined by the trained Go-NoGo geometry predicts choice probabilities

We next asked whether better geometry was synonymous with better performance. Individual capacity and D’s correlated significantly (**fig. 6c-d**, Spearman *ρ* = 0.46, p = 3.0 *×* 10^−4^) across Go/NoGo angles, but this was largely due to capacity tracking the Go/NoGo angle, and it could not predict performance at fixed task difficulty (**fig. 6e**, partial Spearman *ρ* = −0.05, p = 0.72). Therefore, the magnitude of manifold capacity did not by itself predict individual performance. However, this independence was not absolute across measures since a scale-normalized index of Go/NoGo separation in the population PaCMAP embedding retained a modest within-difficulty correlation with performance (**fig. S12**, partial Spearman *ρ* = 0.30, p = 0.03). This weak trial-averaged relationship is expected even if the geometry serves behavior. Both measures are indirect, and performance ultimately depends on how well downstream circuits read V1 out rather than on the separability available in it (see Discussion). We therefore asked the more direct question of whether the position of individual trials within the learned geometry predicts the animals’ choices.

We have shown so far that training induces a geometric reshaping that strengthens and uniformizes the spatial structure of the Go and NoGo representations. Importantly, we found that multiple measures (trajectory angle, centroid distance, capacity, SVMs) show persistent values at the three first Go/NoGo angles in Trained animals, while they track the angle’s decrease in Naive animals. We therefore hypothesized that during training the geometry is constrained such as the trajectories of the task cue representations consistently occupy a location of the state space that can be read out by downstream targets. A way to test this hypothesis would be to determine whether the position of single trials in the space defined by the trained, disentangled geometry is predictive of choice probability. More specifically, we would predict that the animal’s error probability would be greater if the neuronal activity during the trial was away from the centroids of the defined task cues’ object manifolds. Since animals have a licking bias (lick first, then decide) and are reliably licking for Go trials (**fig. 1d**), we focused on NoGo trials, and computed their distance to the Go point cloud centroid by projecting them on the Go-NoGo centroid axis in the PaCMAP space (**fig. 6f**). We then computed the relationship between this position (How Go-like is the NoGo trial in the population’s geometry?) and the licking probability (How likely is the animal to choose Go in those trials?), and binning trials along this axis (bin width 20, sliding window step 5 trials). If our hypothesis is true, then there should be a negative relationship between distance to Go centroid and licking probability. Different animals exhibited different distance ranges, as well as different licking ranges (**fig. 6g & S13)**. To compare the relationships across animals, we normalized the distances by bin ranks, and normalized the licking by z-scoring it within each session (**fig. 6h**). We then computed the slopes of the distance to Go vs. licking probability relationships for the whole cohort (n = 60). The majority of these slopes were negative, significantly more than expected by chance (**fig. 6i**, 47/60, binomial test p = 6.1 *×* 10^−6^), the median slope was −1.99 and sat well outside the null distribution of median slopes obtained by shuffling the licking responses across trials (p < 5 *×* 10^−4^). This result was similar when conducting the same analysis in the native space without dimensionality reduction (**fig. S13**, median slope = −2.01, binomial test p = 3.8 *×* 10^−7^, median vs null p < 5 *×* 10^−4^). Therefore, in NoGo trials, neuronal trajectories outside the NoGo object manifold and towards that of the Go were more error prone. Although its overall separability does not by itself set the level of behavioral performance, the trial-by-trial position of a stimulus within this geometry is reflected in the animal’s choices, consistent with the reorganized V1 representation being read out during the decision.

## Discussion

### Summary of the results

Training for a task is known to induce changes in the neuronal population response to the task cues. Here, we show that training in an orientation discrimination task reorganizes the geometry of stimuli representations at the earliest stage of cortical visual processing. Comparing V1 population responses in mice performing an orientation discrimination task to Naive mice viewing the same stimuli, we found that training reshaped the Go and NoGo object manifolds such that it followed the theoretical predictions for improved disentanglement and linear separability. Indeed, the two manifolds in mice performing the task were more compact, had lower dimensionality, and were pulled further apart in the neural state space than in Naive mice viewing the same set of stimuli. These changes were paired with their predicted functional consequence since both manifold capacity and linear classification accuracy were improved, indicating a higher separability of the task cues in trained animals. Importantly, training-associated changes in the manifolds can be separated in two categories: static changes (radius and dimensionality compression, orthogonalized trajectories) that were constant during the whole duration of the stimulus representation, and dynamic changes (centroid distance and separability) that steadily increased during the trial. We demonstrated that none of those changes in the geometry could be attributed to the difference in behavioral engagement; the geometric reshaping unfolded before and extended during decisions in the task. The learned geometry settled a Go/NoGo relationship in the neural state space that could be linked to the animals decision, since the position of a trial on the Go/NoGo centroid axis was predictive of licking probability, consistent with contributing to the decision process.

### Read out mechanism

Our results dissociate two ways geometry could matter for behavior, based on a dissociation we observed: the overall amount of separability did not predict performance within a difficulty level, but the position of individual trials within the learned geometry did. Part of this dissociation is expected from measurement alone, because task D^*′*^ underestimates the animals’ actual capability, which probe conditions reveal to exceed expressed performance (36), and capacity is estimated from manifolds sampled from a small fraction of the relevant population. But the dissociation also follows from the architecture our data suggest. The animals were trained extensively on an easy version of the discrimination task whose representations are readily separable. In this regime, performance is limited less by the separability available in V1 than by how well downstream circuits have learned to read it out, and mice are known to leverage the information available in V1 suboptimally (22, 23). Our trial-level result offers a glimpse of how this readout operates. Concretely, what reaches the decision is not a summary of the representation’s quality, but the position of each response within the learned geometry. We propose that a downstream integrator accumulates evidence from V1 by reading the activity coming from specific positions of the population state space (37). Because the learned manifolds are linearly separable, this readout requires only a weighted sum of V1 activity (4), with training having adjusted those weights (38, 39) so that responses falling at the learned manifold locations drive licking or its suppression.

In that hypothesis, V1 outputs a graded probability that the stimulus is the Go or the NoGo, which downstream circuits accumulate and threshold into behavior with added noise, consonant with drift-diffusion accounts of decision making (35). This mechanism aligns with multiple other aspects of our results. The active maintenance of the manifold distances and locations throughout the trial, would provide the sustained evidence required by such an integrator. Moreover, a readout of this kind constrains its own inputs: once the weights have been learned, its performance depends on the same neurons, or locations in state space. Keeping the learned geometry stable is therefore advantageous, and we observed the trained Go/NoGo configuration to exhibit such stability. The trajectory angle, centroid distance, capacity and classification accuracy were more constant across days than in Naive animals where the geometry tracked the Go/NoGo angle. Neurons were not tracked across sessions, so we cannot confirm that the same cells carry this configuration across days. Nevertheless, the stereotypy of the geometry is compatible with a representation held stable for the benefit of a trained readout, as if each new NoGo stimulus were mapped onto a pre-established scaffold rather than represented anew.

### Feature-encoding link & interpretability

This stereo-typed mapping is precisely what we observed at the feature-encoding level in this paradigm: as the NoGo orientation stepped closer to the Go (from 90° to 65°), the evoked activity kept concentrating at the same orientation encoding domain of 90°-preferring neurons instead of following the presented orientation (22). Additionally, training promoted more accurate orientation representations with reduced trial-to-trial noise by consistently recruiting the same task-relevant neurons (20), the feature-space counterpart of the compact, stereotyped manifolds we describe here. The correspondence also lives in the probabilistic encoding: the relative activation of the learned Go and NoGo domains predicts the animals’ decision probabilities (22), a code that the position of single trials along the Go–NoGo centroid axis recapitulates at the geometric level. This mapping between levels of description matters beyond our own results. As the field increasingly adopts abstractive geometric characterizations of population activity, there is a risk of obfuscating the underlying physiology. Here, each geometric change can be linked to interpretable feature-encoding changes. The changes reported in V1 after training on similar tasks are diverse, and sometimes opposite at the single-cell level: increases in response amplitude (19, 40), in the number of responsive neurons (17, 19) and in their selectivity (16–19), but also response suppression producing sparser codes (20, 21). These are different routes to the same computational end, a better signal-to-noise ratio leading to a wider separation of the task representations. All register in the same way in the manifolds’ radius, distance and dimensionality, suggesting that geometric measures tap into the core principle these diverse implementations realise, while remaining interpretable precisely because they can be anchored to the feature-level changes that produce them.

### Linearity of the geometric measures

Manifold capacity is, by construction, a linear measure, which asks how separable the Go and NoGo manifolds are to a linear readout, aligned with the object-manifold view in which learning untangles representations into linearly separable form (4, 6, 9). A related but slightly different strand of population-geometry research instead foregrounds the low-dimensional dynamics along which activity evolves and the often nonlinear structure of the underlying manifold (41–44). These are complementary emphases rather than competing theories (7), differing mainly in whether they foreground the separability of representations at each moment or the temporal flow linking them. Our analysis engages both, since by tracking separability across the trial, we characterise the reorganisation as a temporally extended process rather than a static repositioning of response clouds. It is linear in a second, deeper sense as well capacity and the readout axis treat the task structure as linearly separable - yet orientation is a circular variable whose V1 representation is intrinsically curved, making the discrimination of nearby orientations a locally non-linear problem. Learning may therefore reshape this nonlinear structure rather than simply pulling flat manifolds apart, and in this system it demonstrably does, because after training, V1 warps the orientation space into discrete, task-defined categories, such that cues flanking the trained orientations come to be represented more similarly and the Go and NoGo stimuli engage distinct representational domains (20, 22). Our linear geometric changes (i.e., larger centroid distance, smaller radius) may thus be the population-level projection of this categorical warping of the underlying feature space. However, because linear separability is a lower bound on what a nonlinear readout could recover, our measures may understate the reorganisation. Characterising this nonlinear structure directly, and relating it to the categorical warping of the orientation space, is an important direction for future work.

### V1 as a locus of geometric reorganization

To our knowledge, this study provides the first demonstration that learning reshapes the geometry of visual stimulus manifolds in the primary visual cortex. Comparable disentangling had so far been reported downstream, in parietal, temporal and motor areas (3, 8, 10, 11). Finding such geometric changes at the earliest cortical stage is consistent with the architecture of the visual system, which operates as a set of nested loops rather than a sequence of relays. Each area projects forward while receiving dense feedback from downstream areas that refine the very representation they read from, as when feed-back from MT sharpens motion and figure-ground signals in V1 (45–47). That way, improving an early representation improves the evidence available to every subsequent processing stage. In this scheme, V1 acts as a high-resolution workspace onto which higher areas write intermediate results and from which they read refined ones (13). We propose that the reorganization we observe serves exactly this role: the decision is not realised in V1, but by pulling the Go and NoGo manifolds apart, compressing them, and holding them separable across the trial, V1 supplies the downstream readout with better-separated evidence. This V1 involvement may be especially prominent in the mouse, whose visual system is shallower and whose V1 receives proportionally more extensive top-down projections than that of primates (48, 49), potentially placing a relatively larger share of the computation in V1 itself.

### Two regimes of geometric change: static and dynamic

The reorganization we observe is not monolithic, but separates into two components with distinct temporal profiles. A static component, the compaction of the manifolds through reduced radius and dimensionality, was present from the earliest evoked frames and held roughly constant throughout stimulus presentation. A dynamic component, namely the separation of the manifolds through increased centroid distance, manifold-capacity and SVM accuracy, emerged only after the initial transient and was actively amplified across the trial. The two follow different time courses, and compaction can occur without a separability gain, as when sequence learning sparsens and decorrelates V1 population activity yet leaves stimulus decoding unchanged (50). This indicates that they are not one process observed twice, but two dissociable arms of the reorganization.

The static component is consistent with the learning-related changes discussed above, i.e. sharper, sparser, more consistently recruited representations. Changes expressed from response onset can be produced by plasticity local to V1 or its inputs, through refinements of feedforward tuning and of horizontal and recurrent connectivity that reshape the evoked response as it is generated (51, 52), but can also be produced by top-down tonic signals like state and context (53, 54). Because such mechanisms interact with the feedforward sweep itself, their geometric signature is expected from the first frames, as we observe.

The dynamic component is less well characterized, partially because taking an average across timepoints has been the most common approach in the field. Learning-related reshaping of V1 population responses has been reported, including the orthogonalization of stimulus responses (21) and the untangling of sequence representations in principal-component space (50), but the progressive, within-trial construction of manifold separability, as opposed to a static difference, has not been described yet in V1. Its late emergence and continued amplification place it in the epoch shaped by recurrent and cortico-cortical feedback computation rather than the feedforward sweep (55, 56). This fits causal evidence that V1’s contribution to perception extends beyond the feedforward sweep: figure-ground perception requires V1 activity extending beyond it, and longer for harder discriminations (47), perturbing the late phase of the V1 response through its parvalbumin interneurons impairs discrimination even when the early sweep is spared (57), and the window during which V1 is causally required extends as task demands grow (58). The dynamic effect is also compatible with the trial-to-trial variability of the learned population transformation reported in mouse V1, which was attributed to an active circuit mechanism rather than fixed synaptic change (21).

We therefore hypothesize that the two components have distinct mechanistic origins, an early, locally or feedback-driven reshaping of the evoked representation and a late, recurrently driven sharpening of its separability.

### Limitations and future directions

Our study’s limitations leave important questions open. First, it compares snapshots of two end states, expert animals and naive ones, and thus captures the outcome of the geometric reorganization rather than its elaboration. Understanding how this geometry is built will require following the same neuronal populations longitudinally throughout training. Beyond establishing causality of learning in the reorganization, such recordings would reveal whether the static and dynamic components emerge together or follow different timelines, directly testing the distinct mechanistic origins we discussed above. Second, our account stops at V1’s output, but if V1 prepares a refined, more separable representation that is read out during the decision, then the key open questions lie downstream. Which areas provide the feedback that shapes the refinement, where is this signal read out, and how is it transformed as it propagates toward decision-related structures are natural follow-up questions. Tracking the fate of this representation across the hierarchy, together with the trajectory of its construction during learning, would turn the geometric description we provide here into a mechanistic account of how the early sensory cortex comes to serve the decision process.

## Materials and Methods

### Animals and surgery

Adult (3–6 months old) male and female mice were used, housed on a 7:00–19:00 light/dark cycle (humidity 30–70%, temperature 68–72 °F). The Trained cohort comprised 10 C57BL/6 mice (Jackson Laboratories #000664) in which the fluorescent calcium indicator GCaMP6f was expressed by viral injection of AAV1.eSyn.GCaMP6f.WPRE.SV40 (UPenn Vector Core). The Naive cohort comprised 7 Ai163 *×* Camk2a-Cre transgenic mice (Jackson stock #005359) expressing GCaMP6s in excitatory cells, with no viral injection. Ten minutes after systemic analgesia (carprofen, 5 mg per kg body weight), mice were anaesthetised with isoflurane (5% induction, 1.2% maintenance) and placed in a stereotaxic frame. Body temperature was maintained at 37 °C, pressure points and incision sites were infiltrated with 2% lidocaine, and the eyes were protected with artificial-tear ointment. A custom lightweight metal head-bar was fixed to the skull with Vetbond (3M), and a water-retaining recording chamber for the water-immersion objective was built with dental cement (Ortho-Jet, Lang). A circular craniotomy (3 mm diameter) was made over the primary visual cortex (V1). In the trained cohort, AAV1.eSyn.GCaMP6f.WPRE.SV40 (UPenn Vector Core) was injected at three sites separated by 500 µm around the centre of V1 (stereotaxic coordinates −4.0 mm AP, +2.2 mm ML from bregma) with a MicroSyringe Pump Controller Micro 4 (World Precision Instruments) at 30 nL/min; injection began 550 µm below the pial surface and the pipette was raised in 100 µm steps to 200 µm below the dura, delivering 0.7 µL in total across all depths. A 3 mm-diameter cover-with better-separated evidence. This V1 involvement may be especially prominent in the mouse, whose visual system is shallower and whose V1 receives proportionally more extensive top-down projections than that of primates (48, 49), potentially placing a relatively larger share of the computation in V1 itself.

### Two regimes of geometric change: static and dynamic

The reorganization we observe is not monolithic, but separates into two components with distinct temporal profiles. A static component, the compaction of the manifolds through reduced radius and dimensionality, was present from the earliest evoked frames and held roughly constant throughout stimulus presentation. A dynamic component, namely the separation of the manifolds through increased centroid distance, manifold-capacity and SVM accuracy, emerged only after the initial transient and was actively amplified across the trial. The two follow different time courses, and compaction can occur without a separability gain, as when sequence learning sparsens and decorrelates V1 population activity yet leaves stimulus decoding unchanged (50). This indicates that they are not one process observed twice, but two dissociable arms of the reorganization.

The static component is consistent with the learning-related changes discussed above, i.e. sharper, sparser, more consistently recruited representations. Changes expressed from response onset can be produced by plasticity local to V1 or its inputs, through refinements of feedforward tuning and of horizontal and recurrent connectivity that reshape the evoked response as it is generated (51, 52), but can also be produced by top-down tonic signals like state and context (53, 54). Because such mechanisms interact with the feedforward sweep itself, their geometric signature is expected from the first frames, as we observe.

The dynamic component is less well characterized, partially because taking an average across timepoints has been the most common approach in the field. Learning-related reshaping of V1 population responses has been reported, including the orthogonalization of stimulus responses (21) and the untangling of sequence representations in principal-component space (50), but the progressive, within-trial construction of manifold separability, as opposed to a static difference, has not been described yet in V1. Its late emergence and continued amplification place it in the epoch shaped by recurrent and cortico-cortical feedback computation rather than the feedforward sweep (55, 56). This fits causal evidence that V1’s contribution to perception extends beyond the feedforward sweep: figure-ground perception requires V1 activity extending beyond it, and longer for harder discriminations (47), perturbing the late phase of the V1 response through its parvalbumin interneurons impairs discrimination even when the early sweep is spared (57), and the window during which V1 is causally required extends as task demands grow (58). The dynamic effect is also compatible with the trial-to-trial variability of the learned population transformation reported in mouse V1, which was attributed to an active circuit mechanism rather than fixed synaptic change (21).

We therefore hypothesize that the two components have distinct mechanistic origins, an early, locally or feedback-driven reshaping of the evoked representation and a late, recurrently driven sharpening of its separability.

### Limitations and future directions

Our study’s limitations leave important questions open. First, it compares snapshots of two end states, expert animals and naive ones, and thus captures the outcome of the geometric reorganization rather than its elaboration. Understanding how this geometry is built will require following the same neuronal populations longitudinally throughout training. Beyond establishing causality of learning in the reorganization, such recordings would reveal whether the static and dynamic components emerge together or follow different timelines, directly testing the distinct mechanistic origins we discussed above. Second, our account stops at V1’s output, but if V1 prepares a refined, more separable representation that is read out during the decision, then the key open questions lie downstream. Which areas provide the feedback that shapes the refinement, where is this signal read out, and how is it transformed as it propagates toward decision-related structures are natural follow-up questions. Tracking the fate of this representation across the hierarchy, together with the trajectory of its construction during learning, would turn the geometric description we provide here into a mechanistic account of how the early sensory cortex comes to serve the decision process.

## Materials and Methods

### Animals and surgery

Adult (3–6 months old) male and female mice were used, housed on a 7:00–19:00 light/dark cycle (humidity 30–70%, temperature 68–72 °F). The Trained cohort comprised 10 C57BL/6 mice (Jackson Laboratories #000664) in which the fluorescent calcium indicator GCaMP6f was expressed by viral injection of AAV1.eSyn.GCaMP6f.WPRE.SV40 (UPenn Vector Core). The Naive cohort comprised 7 Ai163 *×* Camk2a-Cre transgenic mice (Jackson stock #005359) expressing GCaMP6s in excitatory cells, with no viral injection. Ten minutes after systemic analgesia (carprofen, 5 mg per kg body weight), mice were anaesthetised with isoflurane (5% induction, 1.2% maintenance) and placed in a stereotaxic frame. Body temperature was maintained at 37 °C, pressure points and incision sites were infiltrated with 2% lidocaine, and the eyes were protected with artificial-tear ointment. A custom lightweight metal head-bar was fixed to the skull with Vetbond (3M), and a water-retaining recording chamber for the water-immersion objective was built with dental cement (Ortho-Jet, Lang). A circular craniotomy (3 mm diameter) was made over the primary visual cortex (V1). In the trained cohort, AAV1.eSyn.GCaMP6f.WPRE.SV40 (UPenn Vector Core) was injected at three sites separated by 500 µm around the centre of V1 (stereotaxic coordinates −4.0 mm AP, +2.2 mm ML from bregma) with a MicroSyringe Pump Controller Micro 4 (World Precision Instruments) at 30 nL/min; injection began 550 µm below the pial surface and the pipette was raised in 100 µm steps to 200 µm below the dura, delivering 0.7 µL in total across all depths. A 3 mm-diameter coverslip was then placed over the dura, flush with the skull surface, and secured with Vetbond and dental cement. Amoxicillin (0.25 mg/mL) was provided in the drinking water for the 5 days following surgery, and mice recovered for at least 3 weeks to allow gene expression. Ten mice were trained on the discrimination task and imaged during performance. Seven Naive mice passively viewed the identical stimulus sequence, yielding 102 imaging sessions in total.

### Behavioural task and training

Mice were water-deprived to 85% of their body weight and acclimated to head fixation on a spherical treadmill in a custom-built, sound-proofed rig equipped with a monitor and an infrared-beam lickometer. Data-acquisition boards (National Instruments, Arduino) actuated the reward solenoids and recorded licking under a custom MATLAB program. Mice were trained on a Go/NoGo orientation-discrimination task. Drifting sine-wave gratings oriented 45° below the horizontal were paired with a water reward and the animal was expected to lick (Go trial); gratings orthogonal to the Go signal (135°) signalled the absence of reward and the animal was expected to withhold licking (NoGo trial). Gratings were presented full-screen and shared the same temporal frequency (2 Hz), spatial frequency (0.04 cycle per degree) and duration (3s); contrast was 75% during training and 25% during recording. Water was delivered if the mouse licked during the third second of a Go stimulus. The inter-trial interval was 3 s after correct trials (Hit or Correct Rejection) and increased to 6s after incorrect trials (Miss or False Alarm) as negative reinforcement. Performance was quantified with the sensitivity index D^*′*^ = −Φ-1(Hit rate) Φ-1(False-alarm rate), where Φ-1 is the inverse normal CDF; animals were considered expert when training performance exceeded D^*′*^ = 1.7. As a non-discrimination reference we computed the “best random D^*′*^” ceiling (as in Corbo et al., 2025), the sensitivity of a biased-but-non-discriminating strategy at a favourable operating point (the 77.5th/22.5th-percentile (p=0.2252 = 0.05) of the Hit/FA distribution over the typical trial count, n = 128), giving D^*′*^ = 1.68.

During imaging, the Go orientation was fixed at 45° while the NoGo orientation was set to 135°, 90°, 75°, 70°, 65° and 60° across six consecutive days, so that the Go/NoGo angular separation decreased from 90° to 15° (Day 1: 90°, Day 2: 45°, Day 3: 30°, Day 4: 25°, Day 5: 20°, Day 6: 15°). Recording sessions were limited to 120 trials to keep mice motivated. Each imaging session included an orientation-tuning block of passively viewed drifting gratings (12 orientations evenly spaced by 30°, contrast 100%, 1.5 s each) used to estimate each neuron’s preferred orientation.

### Two-photon imaging and preprocessing

Functional imaging was performed at 15 frames per second on a resonant-scanning two-photon microscope (Neurolabware) powered by a Ti:Sapphire Ultra-2 laser (Coherent) set to 910 nm; the beam was focused 200 µm below the cortical surface through a 16×, 0.8 NA Nikon water-immersion objective, at laser power below 70 mW. Frames (512 *×* 796 pixels) were acquired with Scanbox (Neurolabware) and realigned offline with the Scanbox motion-correction algorithm. Regions of interest were segmented automatically with Suite2p. Neuropil contamination of the somatic fluorescence was removed by subtracting the mean fluorescence of a 2–5 µm ring surrounding each ROI (excluding the somata of neighbouring neurons) and then adding back the across-time median of the subtracted background. Fractional fluorescence dF/F = (F − F_0_)/F_0_ was computed with F_0_ the median raw fluorescence during every inter-trial interval, and deconvolved into action-potential-related events (APrEs) with MLspike (24). Neurons were included when the correlation between the measured dF/F and the dF/F reconstructed from the inferred APrEs exceeded 0.8, together with Suite2p “iscell” > 0.8, ROI area > 15 µm^2^, roundness (the percentage overlap with a circle centred on the ROI) > 20%, and a response probability at the preferred orientation (tuning block) > 0.1. The orientation-tuning block was excluded from task analyses.

For time-resolved population analyses (frame period 64.6 ms; 140-frame trials with 1.5 s of pre-stimulus baseline), each trial was represented in consecutive non-overlapping windows of 3 frames (textasciitilde190 ms; 3-frame step), each labelled by the time of its last frame, spanning −1 to 4 s relative to stimulus onset. Windows were labelled by their last frame and time-courses are plotted against that time. Within-window APrE counts were summed and each neuron was z-scored to its own pre-stimulus (t < 0) window distribution, with the per-neuron standard deviation floored at 0.12 (integrated-APrE units - see fig. S14). The floor is a reliability cap: because the pre-stimulus windows are sparse integer spike counts, the standard deviation is estimated from few events for quiet neurons, and dividing by an unreliably small value over-amplifies them.

### Population response magnitude and sparsening. Population-mean activity

Population activity was summarised as the mean across neurons of the z-scored, window-integrated APrE signal, computed separately for Go and NoGo trials and each session and then averaged across sessions (mean *±* SEM; trained n = 60, naive n = 42 sessions per stimulus). Naive-versus-trained differences (separately for Go and NoGo) were tested with the cluster-based permutation test (Statistics), and Go-versus-NoGo differences (within each group) with its paired, sign-flip variant.

#### Fraction of responsive neurons

Responsive neurons were characterized using a binomial test. For each neuron and trial, the windowed APrE count was binarized (≥ 1 APrE = active). Each neurons’ baseline probability was computed from the 1s immediately preceding stimulus, as well as the associated Clopper–Pearson 95% upper confidence bound (*α* = 0.05). In each of three post-stimulus windows (0–1, 1–2, 2–3 s) a neuron was scored responsive if its activation probability exceeded this baseline bound, and the session-level metric was the fraction of responsive neurons. For the main comparison Go and NoGo trials were pooled (“all task trials”).

### Manifold visualization

Per-session activity was embedded in three dimensions with PaCMAP (25) (n_neighbors = 10, MN-ratio = 0.5, FP-ratio = 2, PCA initialization, version 0.8.0). Two normalizations were applied before embedding. Each trial’s pre-stimulus mean (−1.5 to 0 s) was subtracted per neuron, so that the embedding reflects each trial’s evolution from its own baseline rather than absolute baseline differences between trials. Each neuron was then z-scored by its standard deviation over the whole session (all trials and time windows), placing neurons on a comparable scale for the embedding’s distance metric. In each session, per-window point cloud centroids were computed separately for Go and NoGo trials to give the two centroid trajectories.

Because neurons are not tracked across sessions, cohort figures were brought into a common frame by a two-pass Procrustes registration of the stacked Go and NoGo centroid trajectories (rotation and reflection, no scaling). Each session was aligned to the medoid session of its recording day (the session with the smallest summed centroid-trajectory distance to the others of that day), and each day’s medoid was then aligned to day 1. A single viewing angle, computed from the day-1 average trajectories, was applied to all cohort panels.

To summarize the configuration across days, the aligned Go and NoGo clouds were projected onto a common twodimensional discriminant frame (axis 1, the pooled NoGo minus Go mean-difference vector; axis 2, the leading principal component of the residual), and the 1-SD covariance ellipse of each stimulus was overlaid across days.

#### Trajectory angles

For each session we computed the angle *θ* between the Go and NoGo centroid-trajectory displacement vectors over the early response, between 0 and 0.8 s after stimulus onset. Because *θ* is an angle between two vectors defined within the same session, it is invariant to the arbitrary per-session embedding frame and requires no cross-session alignment.

#### Cross-method validation

The same activity was embedded with four additional methods (UMATO, classical MDS, Isomap and non-metric MDS) and the Go and NoGo clouds compared in the same discriminant projection with 1-SD co-variance ellipses. For this comparison neurons were baseline z-scored rather than globally z-scored. Isomap and UMATO short-circuit under a global z-score: that metric weights each neuron by the inverse of its total variance, down-weighting discriminative neurons whose variance is inflated by their evoked response, so nearest-neighbour graphs acquire cross-cluster edges. A baseline z-score sets each neuron’s scale from the pre-stimulus window and preserves local cluster separability; the global-distance methods (classical and non-metric MDS) and PaCMAP, which applies a PCA pre-projection, are insensitive to this choice.

### Population geometry in the native space. Geometry (centroid distance, radius)

Population geometry was quantified directly in the native state space, without dimensionality reduction. For each session and time window, single trials were represented as population response vectors. For each stimulus (Go, NoGo) we computed the class centroid (the mean response vector) and the class radius (the mean Euclidean distance of single trials to their class centroid), where the centroid distance was the Euclidean distance between the Go and NoGo centroids. All distances were divided by the square root of the number of neurons in the session, so that they are comparable across sessions of different population sizes.

#### Dimensionality (participation ratio, k80, tail slope)

For each session, time window and stimulus separately, principal component analysis was applied to the mean-subtracted single-trial population responses (the within-class covariance), computed on balanced random subsamples of up to 250 neurons and 100 trials per class and averaged over 5 independent subsampling rounds per session. From the resulting eigenvalue spectrum *λ* we derived three measures: the participation ratio D_PR = (Σ*λ*)^2^ / Σ*λ*^2^; k80, the number of components needed to reach 80% of the cumulative variance; and the tail slope, the slope of a linear fit to the log-eigenvalue tail over components 10 to 40. The three measures were averaged across Go and NoGo, tracked across the trial time course, and compared between Naive and Trained groups.

### Population separability

Both the SVM accuracy and manifold capacity measures quantify the linear separability of the Go (45°) and NoGo (day-specific orientation) population representations across the trial, from the windowed single-trial responses.

#### SVM decoding of stimulus identity

In each window a linear support-vector machine was trained to classify Go versus NoGo from the population response vector and evaluated by five-fold cross-validation. To make sessions comparable, neurons were randomly subsampled to a common ceiling (up to 250 per session) and trials were balanced between the two classes (up to 100 per class), and the procedure was repeated over five independent cell-and-trial subsamples. Decoding accuracy was the percentage of correctly classified held-out trials, averaged over folds and repeats to give one value per session and window.

#### Manifold classification capacity

We estimated the manifold classification capacity of the Go and NoGo representations as the critical load, in manifolds per neuron, at which the two manifolds cease to be linearly separable, with higher capacity indicating more separable representations.Per-trial population vectors were baseline-subtracted (each neuron’s pre-stimulus mean removed) rather than z-scored. In each window the Go and NoGo single-trial vectors define two object manifolds in the N-neuron space, entered at their native trial counts. We estimated capacity directly, using the publicly available implementation of Chung, Lee and Sompolinsky (2018), adapted for the two-class Go/NoGo case. In brief, the trial point clouds are projected onto random subspaces of increasing dimension and, at each dimension, a maximum-margin linear classifier tests whether the two manifolds are separable, repeated over many random projections. Capacity was two divided by the smallest subspace dimension at which they are separable in at least half of projections. We used this direct, simulation-based estimate rather than the analytic mean-field expression, which in its regime of validity predicts the same quantity from a manifold’s radius, dimensionality, and correlation structure (Chung, Lee & Sompolinsky, 2018; Chung & Abbott, 2021). This is a specific design choice, because the mean-field theory is derived for many manifolds in generic position and factors out the arrangement of manifold centers, whereas with only two manifolds the separation between the Go and NoGo centers is itself central to the effect. Estimating separability directly keeps this between-class geometry in play, and because the estimate does not draw on our radius, distance, or dimensionality measures, its agreement with them is convergent rather than circular evidence.

### Behavioural-state controls on manifold capacity

Behavioral-state control analyses were performed within the trained cohort, and capacity was estimated as previously described after restricting the trials entering each estimate to a behaviourally defined subset.

#### Licking tertiles

For each trial, licking was summarised by an activity index equal to the integral of the lick trace, z-scored to its own within-session distribution, over the response window 1 to 2 s after stimulus onset. Within each stimulus class separately, trials were divided into low, middle and high tertiles. The capacity time courses were recomputed within the low tertile and within the high tertile, each from the Go and NoGo trials of that tertile, and the two were compared paired within session across the trial. The high-minus-low difference in the window-averaged capacity is reported as a percentage change.

#### Locomotion and pupil size

Locomotion (treadmill speed) and arousal (pupil area) were analysed identically to licking. Each trace was resampled to the imaging frame rate, low-pass filtered (locomotion 1 Hz, pupil 2 Hz), averaged over the analysis window, and z-scored within session, then split into within-class tertiles, and capacity was compared between the high and low tertiles against the count-matched reference. For pupil size, two windows were analysed: a pre-stimulus baseline (−1.5 to 0 s), testing whether the state the animal enters the trial in changes capacity, and a post-stimulus window (1 to 2 s), testing stimulus- and response-linked arousal. Sessions were retained only when the behavioural trace passed a signal-quality check (sufficient within-session variance, with rejection of sessions whose trace was dominated by a stereotyped, trial-locked artifact).

#### Binary licking split

To assess the impact of licking presence/absence (as opposed to licking level), single trials were pooled across all trained sessions and divided into licking and non-licking trials, and capacity was compared between the two at matched trial counts. Because non-licking trials are comparatively rare and capacity is trial-count sensitive, this split is interpreted only relative to the count-matched reference.

#### Neural geometry and behaviour. Geometric divergence onsets and behavioural decision times

For each geometric measure (dimensionality, manifold radius, centroid distance, manifold capacity) the naive-versus-trained divergence onset was taken from the cluster-based permutation test on the per-session timecourses, pooled across days and reported at the centre of the first sustained-significant window. Its uncertainty was estimated by resampling sessions with replacement within each group and repeating the test (1000 bootstrap resamples; 95% percentile interval). Two behavioural decision times were computed per session from the lick traces, which were smoothed (150 ms moving average) and binarised to a lick-on state. The Go/NoGo lick divergence was the first time at which the across-trial Go lick-on probability exceeded the NoGo probability by a sustained margin, defined as the leading edge of the first run over which the difference exceeded 0.10 for at least 0.20s and the lower bound of a trial-resampling confidence interval (300 resamples) stayed above zero, searched over 0 to 6 s. The last correct-rejection lick was the median, across correct NoGo trials, of the time of the last lick within 0 to 2 s. As a description of Go-lick strategy, the strategy timing Got50 was defined per session as the time at which the across-trial Go lick-on probability first reached 50% of its peak above baseline.

#### NoGo axis position and licking

For each session the Go-to-NoGo axis was defined as the unit vector from the Go-trial centroid to the NoGo-trial centroid of the single-trial population representations, at the time window nearest 1.5 s after stimulus onset, and each NoGo trial was assigned its scalar projection onto this axis (its signed position from the Go centroid toward and beyond the NoGo centroid). NoGo trials were sorted by this position and binned in a sliding window of 20 trials stepped by 5, and the lick probability (fraction of trials with a lick response) was computed in each bin. Because the axis scale differs between sessions, each session’s binned lick-probability curve was z-scored and expressed against the within-session near-to-far rank (0 = closest to the Go end, 1 = farthest), and a line was fit over the near 90% of the rank range; its slope quantifies the dependence of licking on axis position, a negative slope indicating that NoGo trials closer to the Go representation have more licking. Significance was assessed at the group level against a null distribution obtained by shuffling the lick outcomes across each session’s NoGo trials and repeating the entire sorting, binning, ranking and slope procedure (2000 permutations), comparing the observed median slope across sessions to the permutation distribution of that median. The analysis was carried out in the PaCMAP space as in fig. 2 and, as a control, directly in the native space.

#### Statistics

All statistical tests were two-sided. Group means were compared with permutation tests, taking the observed difference as significant when it fell outside the 2.5–97.5th-percentile interval of the null distribution obtained by label shuffling. Non-parametric group comparisons used the Wilcoxon rank-sum test for unpaired (between-group) comparisons and the Wilcoxon signed-rank test for paired (within-group) comparisons. Correlations used Spearman or Pearson as noted, and effects of a categorical factor with more than two levels used the Kruskal–Wallis test. Multiple comparisons were controlled with the Benjamini–Hochberg FDR (*α* = 0.05) across the three post-stimulus windows in the comparison of the fraction of responsive neurons (fig. 1). Time-resolved comparisons of the population-mean activity (fig. 1), the native-space geometry and dimensionality measures (fig. 3), the linear-decoding and manifold-capacity timecourses (fig. 4), and the geometric divergence onsets (fig. 6) used a cluster-based permutation test on the per-session timecourses (59). At each timepoint the naive-versus-trained mean difference was compared to a null distribution of 10,000 random label permutations. Contiguous timepoints exceeding the 95th-percentile permutation threshold, together with a minimum effect size of |Hedges’ g| = 0.2, formed candidate clusters, each summarised by its cluster mass (the summed difference within the cluster). A cluster was significant when its mass exceeded the 95th percentile of the null distribution of the maximum cluster mass across permutations, and the reported cluster p-value is the proportion of permutations whose maximum cluster mass equalled or exceeded the observed value (smallest resolvable value 1/10,000 = 1 *×* 10^−4^). For within-group (paired) comparisons the null was formed by randomly sign-flipping each session’s difference timecourse rather than shuffling group labels. The divergence onset is reported at the centre of the first sustained-significant window (the timecourses themselves are plotted against the window’s last frame). Effect sizes are reported as Hedges’ g, with the standardizer matched to the design: the pooled within-condition standard deviation for unpaired (between-group) comparisons, and the average (root-mean-square) of the two within-condition standard deviations for paired (within-group) comparisons. The measure is interpreted against the conventional bands (<0.2 negligible, 0.2–0.4 small, 0.4–0.6 medium, >0.6 large)

## Acknowledgments

This work was funded by The National Institutes of Health National Eye Institute: Grant #R01 EY030860

## Author contributions

J.C. and L.R.C. conceived and designed the study. O.B.E. performed the surgeries and imaging experiments. J.C. and L.R.C. analysed the data. J.C., L.R.C and P-O.P. interpreted the results. J.C. and L.R.C wrote the manuscript with input from P-O.P. P-O.P. supervised the project and acquired funding.

## Data availability

The data supporting the findings of this study will be made publicly available upon publication.

## Code availability

The analysis code used in this study will be made available upon publication.

