## Supplementary figures for "Geometric reshaping of task-relevant representations in the primary visual cortex supports perceptual decisions"

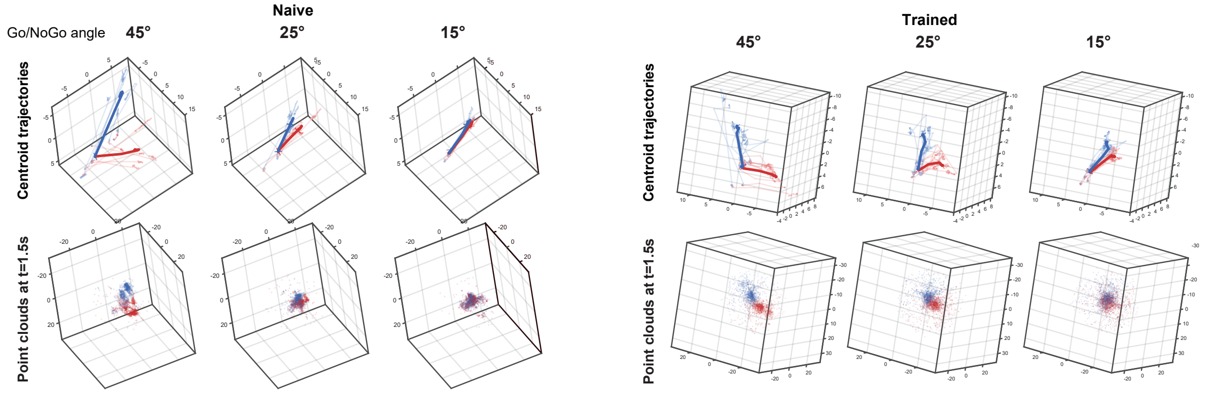


**Figure S1.** Procrustes-aligned trajectories and point clouds for all Naive (n-=7) and Trained (n=10) sessions at Go/NoGo angle of 45°, 25° and 15° respectively from left to right.

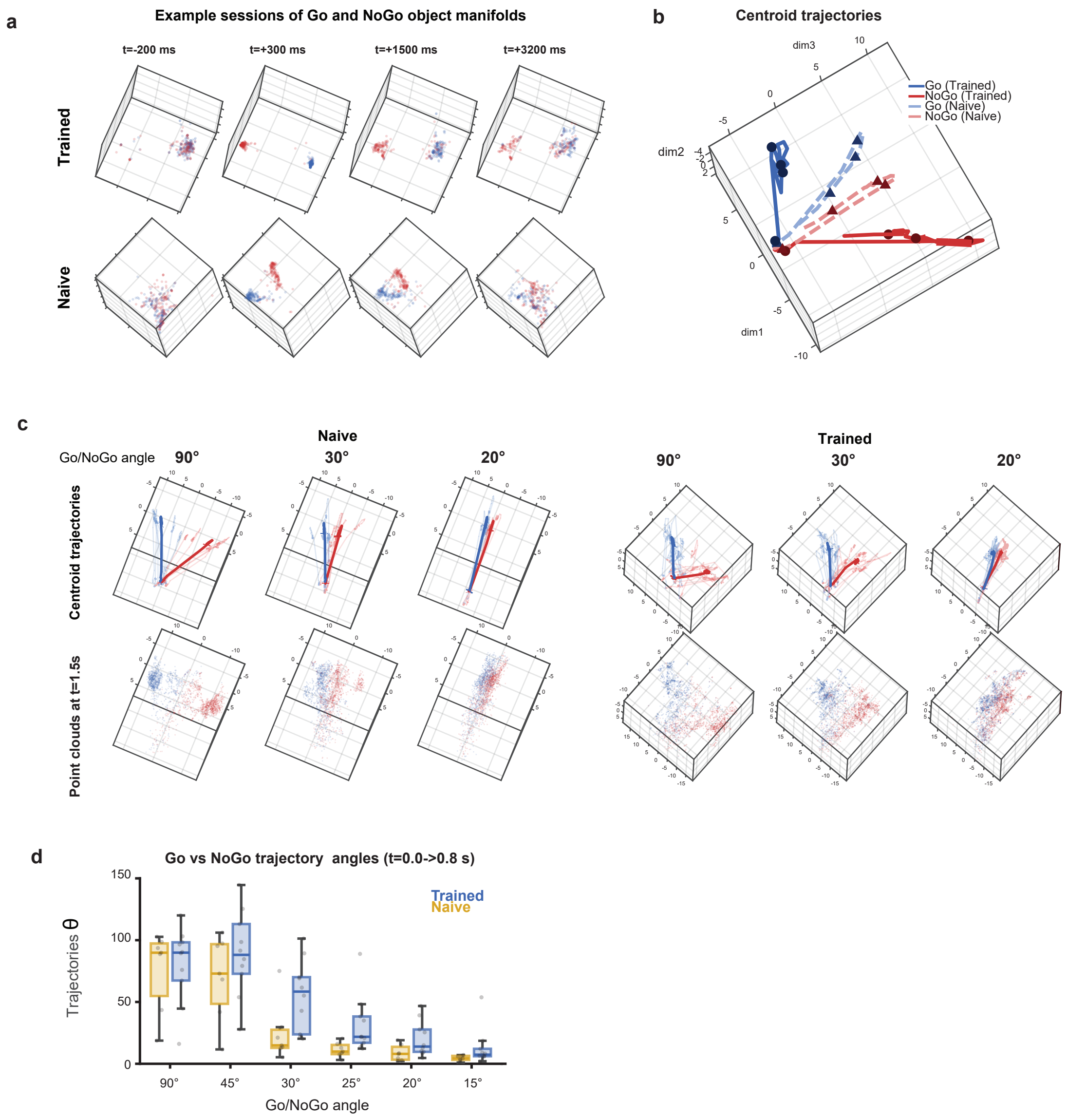


**Figure S2. Visualization of the object manifolds with low-dimensional embeddings with baseline z-score instead of global z-score. a.** PaCMAP projection of the Go (blue) and NoGo (red) single trials point clouds in the neural state space for an example Trained (top) and Naive (bottom) animal recorded at a Go/NoGo angle of 30°. Different snap shots are visualized, from left to right at t=-200ms, +300ms, +1500ms and +3200ms relative to stimulus onset. **b.** Trajectories of the point clouds' centroids from -500ms to 3500ms relative to stimulus onset, for Go and NoGo of the two sessions from a. Darker plain lines indicate the trajectory of the Trained animal's responses, dashed lighter lines that of the Naive animal. Black markers show the time of the snapshots in a. The angle between the Go and NoGo trajectories is indicated as theta. **c.** Procrustes-aligned trajectories and point clouds for all Naive (n-=7) and Trained (n=10) sessions at Go/NoGo angle of 90°, 30° and 20° respectively from left to right. **d.** Trajectory angles summary, computed as the angle between the Go and NoGo point cloud centroid displacement from t=0 to t =800ms in every animal and Go/NoGo angle.

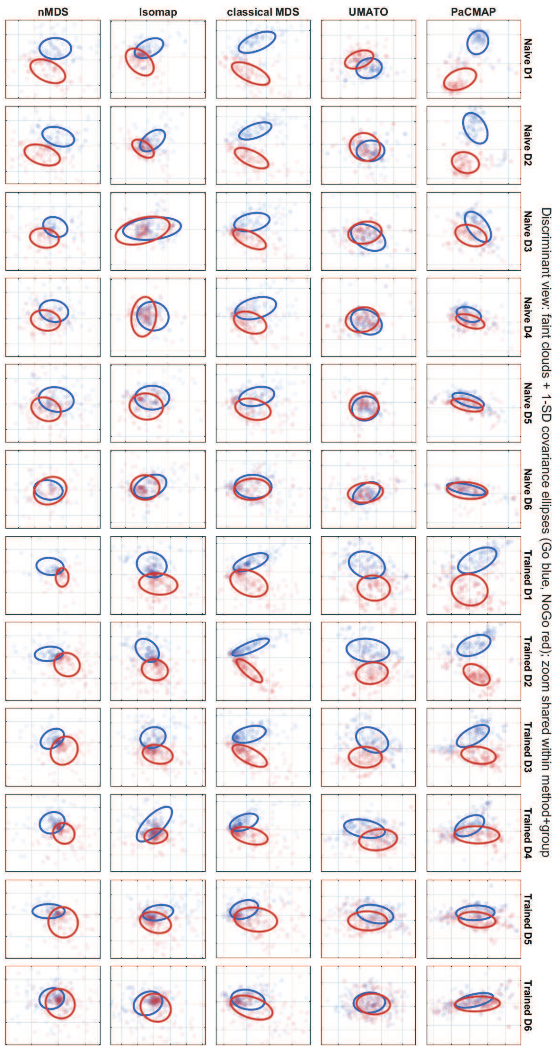


**Figure S3. Comparison of several dimensionality reduction approaches.** Each line is a different method, each column is a different Go/NoGo angle, with the first 6 showing the Naive animals' data, and the 6 last the Trained animals' data. Each panel shows the single-trial Go (blue) and NoGo (red) point clouds in the embedding space, projected onto the most discriminant plane (axis 1: the pooled NoGo minus Go mean-difference vector; axis 2: the leading principal component of the residual variance), with the 1-SD covariance ellipse of each stimulus overlaid.

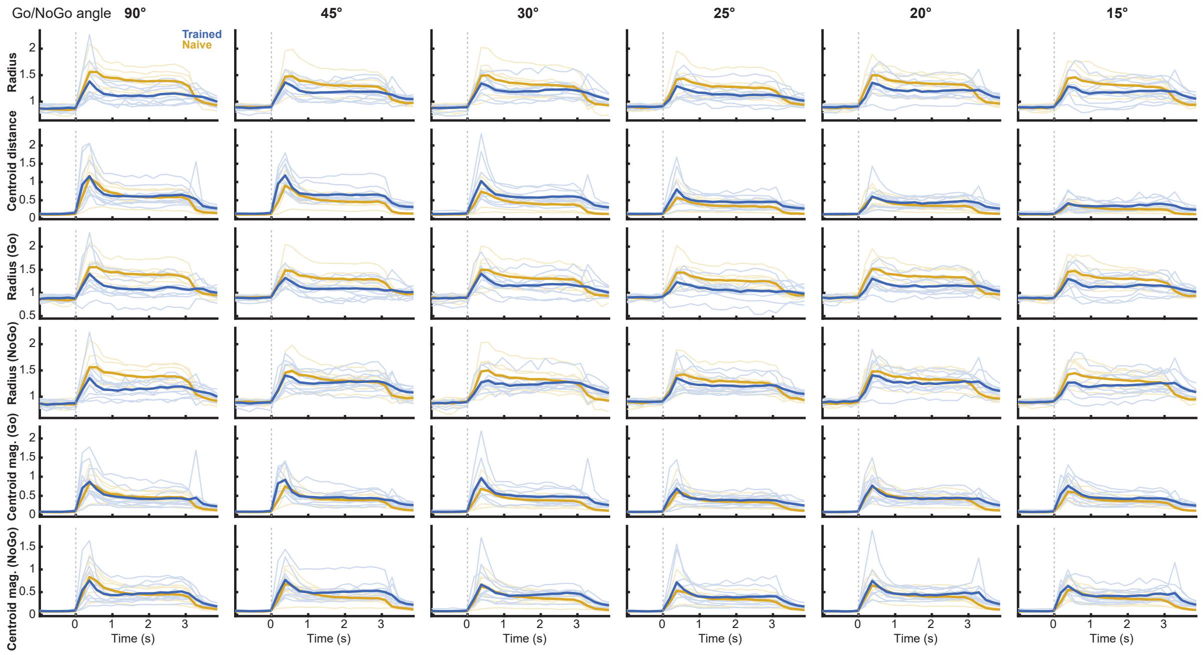


**Figure S4. Time courses of the native-space geometric measures for every Go/NoGo angle.** Each column shows one Go/NoGo angle (90° to 15°) and each row one geometric measure, computed in the native space (no dimensionality reduction) for Trained (blue) and Naive (gold) animals. From top to bottom: the manifold radius (computed as the mean distance of every trial to its within-class centroid, averaged across the two stimulus classes), the Go/NoGo centroid distance (RMS-normalized euclidean distance), the radii of the Go and NoGo point clouds separately, and the magnitude of the Go and NoGo centroids (the distance of each class centroid to the pre-stimulus origin). Thin lines show individual sessions, thick lines show group averages. The dashed line indicates stimulus onset. Note the absence of asymmetry between the radius of the Go and NoGo stimulus representations, and the similar magnitude between Naive and Trained in spite of the stronger activity in Naive, suggesting that the sparseness observed in Trained is a noise reduction.

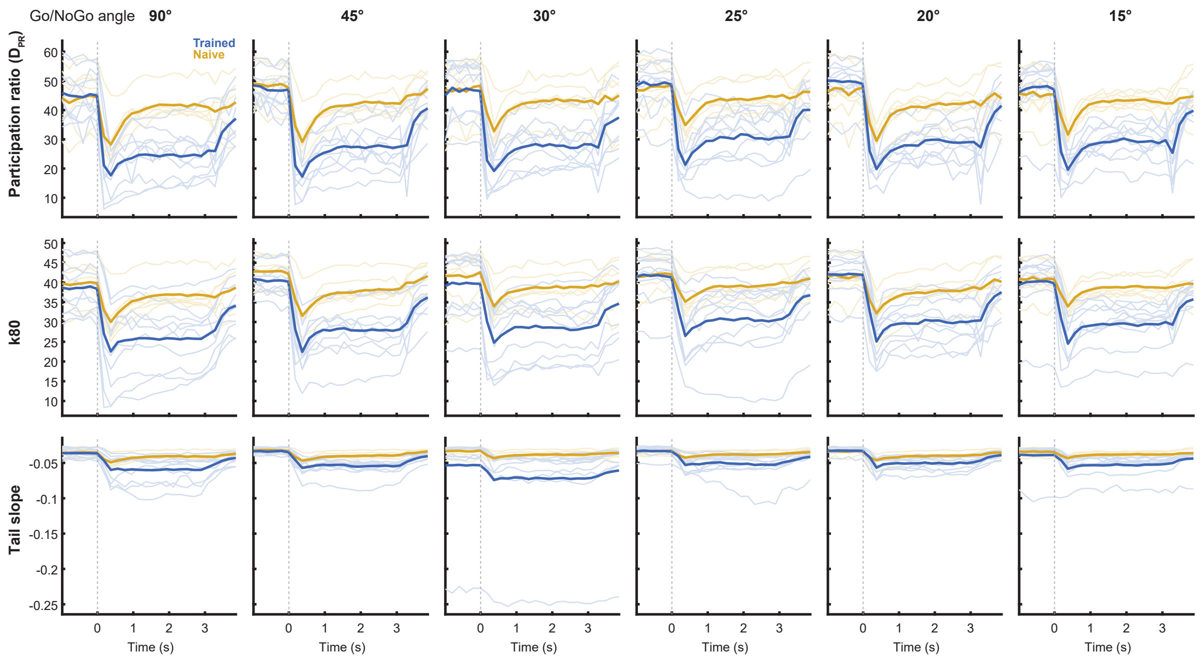


**Figure S5. Time courses of the dimensionality measures for every Go/NoGo angle.** Each column shows one Go/NoGo angle (90° to 15°) and each row one dimensionality measure, computed on the within-class covariance and averaged across the two stimulus classes and subsampling rounds, for Trained (blue) and Naive (gold) animals. From top to bottom: the participation ratio (DPR), k80 (the number of principal components required to explain 80% of the within-class variance), and the slope of the log-eigenvalue spectrum tail. Thin lines show individual sessions, thick lines show group averages. The dashed line indicates stimulus onset.

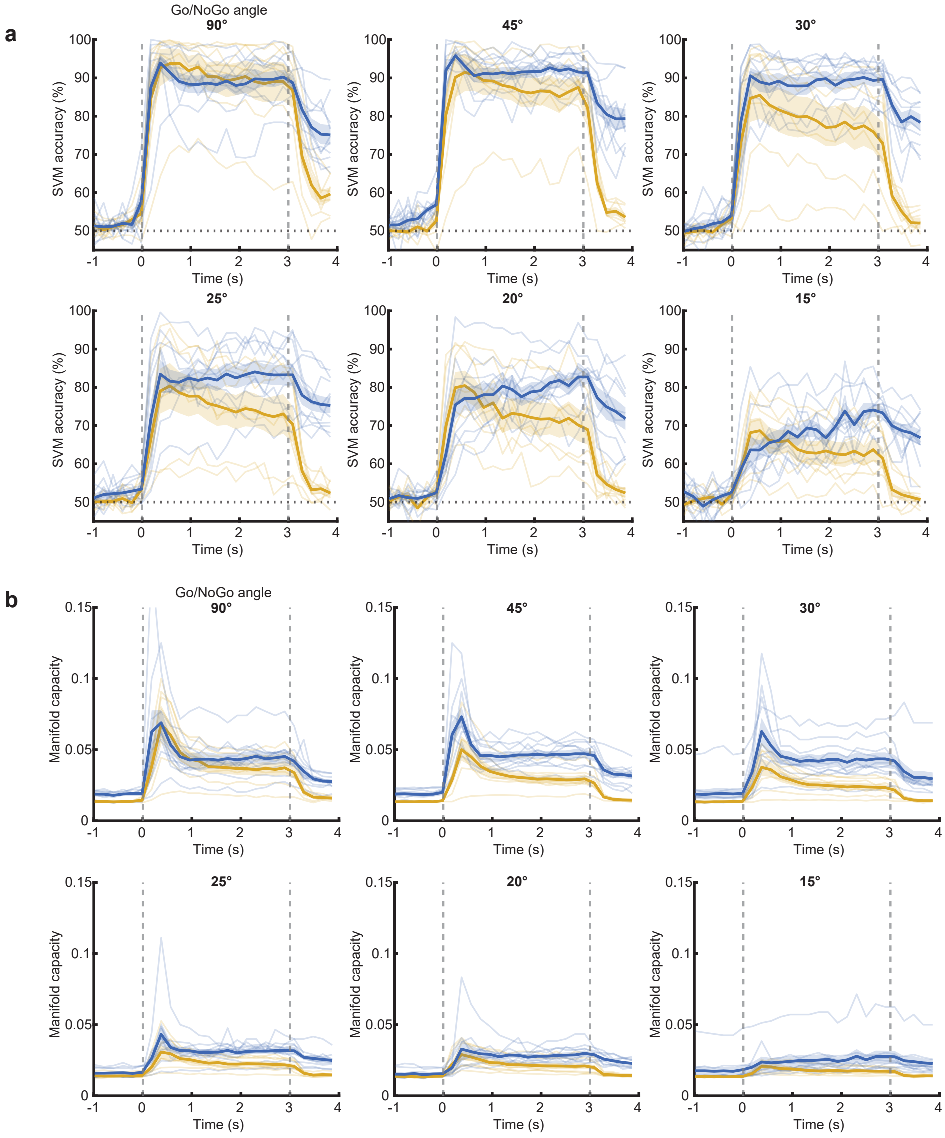


**Figure S6. Time courses of the SVM accuracy and the manifold capacity for every Go/NoGo angle. a.** Go/NoGo classification accuracy time course of linear SVMs trained on Trained (blue) and Naive (gold) population responses, for every Go/NoGo angle (left to right: 90°, 45°, 30°, top row; 25°, 20°, 15°, bottom row). Single sessions are shown as thin lines and group averages as thick lines, with the shaded area indicating s.e.m. Vertical dashed lines indicate stimulus onset and offset, the dotted line the 50% chance level. **b.** Same as a. but for the manifold capacity computed from the Go and NoGo point clouds.

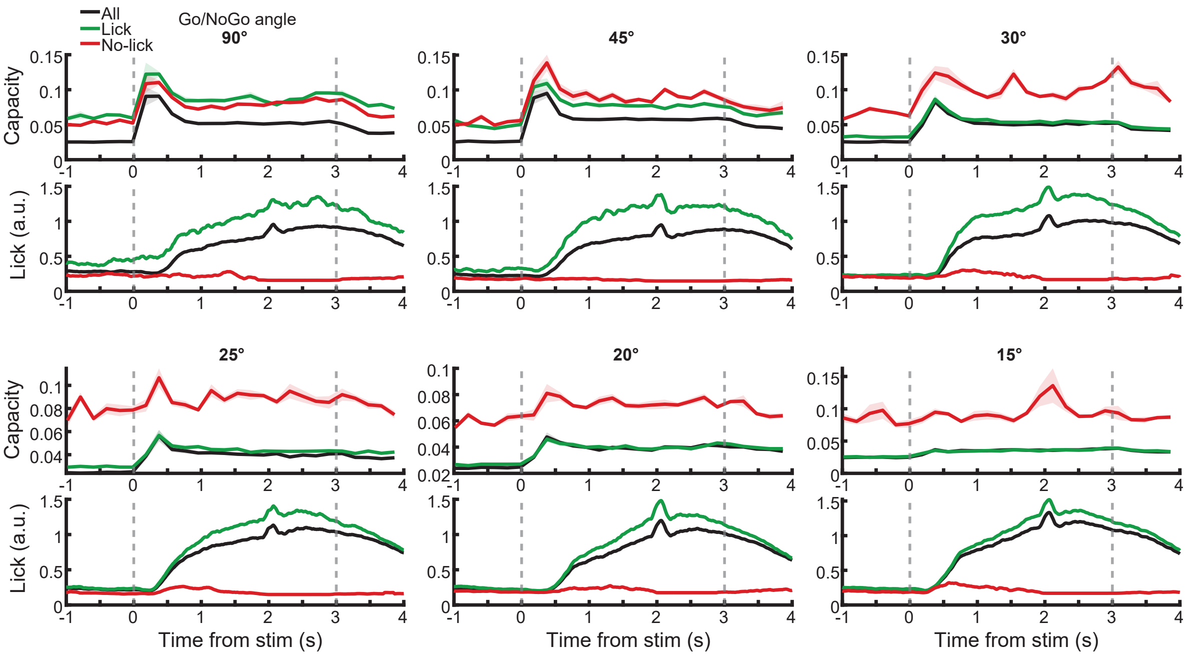


**Figure S7. Capacity computed on a binary lick/no-lick trial split (super-mouse resampling).** For every Go/NoGo angle, manifold capacity is computed from all trials (black), trials with licking (green) and trials without licking (red) in the 2-3s window, in Trained animals. For each Go/NoGo angle, five pseudo-sessions of 500 neurons were assembled by pooling sessions, and 100 pseudo-trials per stimulus class were resampled from each session's eligible trials of the corresponding split, using the same neurons across the three splits. Top rows: capacity time courses (mean ± s.e.m. across pseudo-sessions). Bottom rows: average lick traces reconstructed from the same resampled trials. Vertical dashed lines indicate stimulus onset and offset. Note that although the resampling fixes the number of pseudo-trials, it cannot equalize the number of unique trials available to each split: where a split is scarce, i.e. the no-lick pool at the smallest Go/NoGo angles, the fixed draws repeatedly sample a small set of unique trials, and capacity computed from such under-sampled manifolds is unreliable and inflated, as visible in the elevated no-lick values already present before stimulus onset. Split comparisons are therefore only meaningful where enough unique trials exist in both splits, as at the largest Go/NoGo angles.

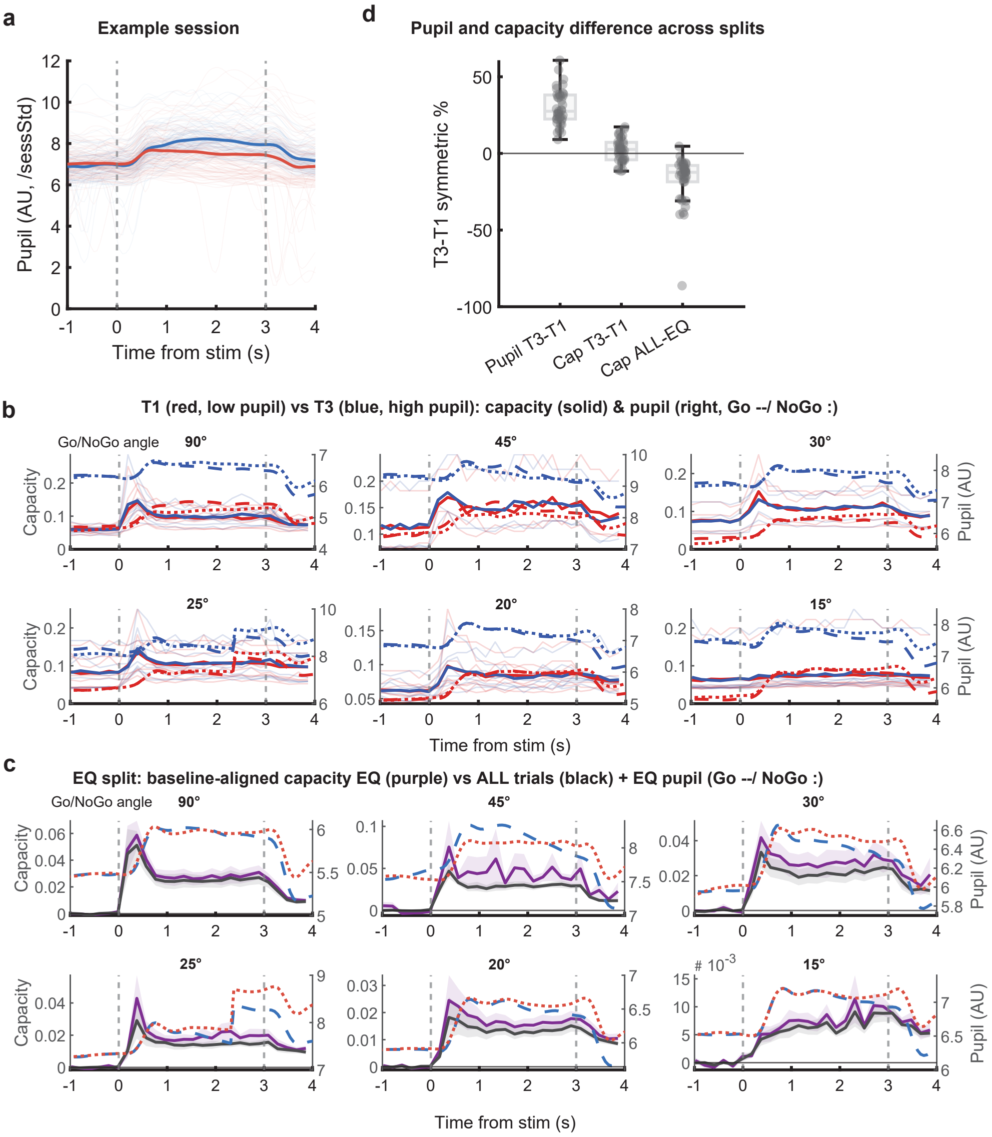


**Figure S8. Pre-trial pupil size does not drive the capacity of the Trained representations. a.** Pupil size time course for an example session. Thin lines show single trials, thick lines the Go (blue) and NoGo (red) averages. Pupil traces were low-pass filtered (2 Hz) and normalized by the session's standard deviation. Dashed lines indicate stimulus onset and offset. **b.** Capacity computed from trials split by pre-stimulus (−1.5–0 s) pupil size into low (T1, red) and high (T3, blue) tertiles, computed within each stimulus class, for every Go/NoGo angle. Left axis: capacity, with thin lines showing individual sessions and thick lines the group averages. Right axis: average pupil traces of the Go (dashed) and NoGo (dotted) trials of each tertile. **c.** Pupil-equalized Go/NoGo control for every Go/NoGo angle. Capacity computed from a subset of Go and NoGo trials matched 1:1 on their pupil size (EQ, purple) compared with the capacity from all trials (black), both baseline-subtracted; shaded areas indicate s.e.m. Right axis: average pupil traces of the matched Go (dashed) and NoGo (dotted) trials, illustrating the removal of the Go/NoGo pupil asymmetry. **d.** Summary of the per-session percentage differences (difference normalized by the mean of the two values): pupil T3−T1 (over the split window), capacity T3−T1 and all-trials versus pupil-equalized capacity (both over the 1–2 s window). Boxes show median and interquartile range, whiskers the range, dots individual sessions.

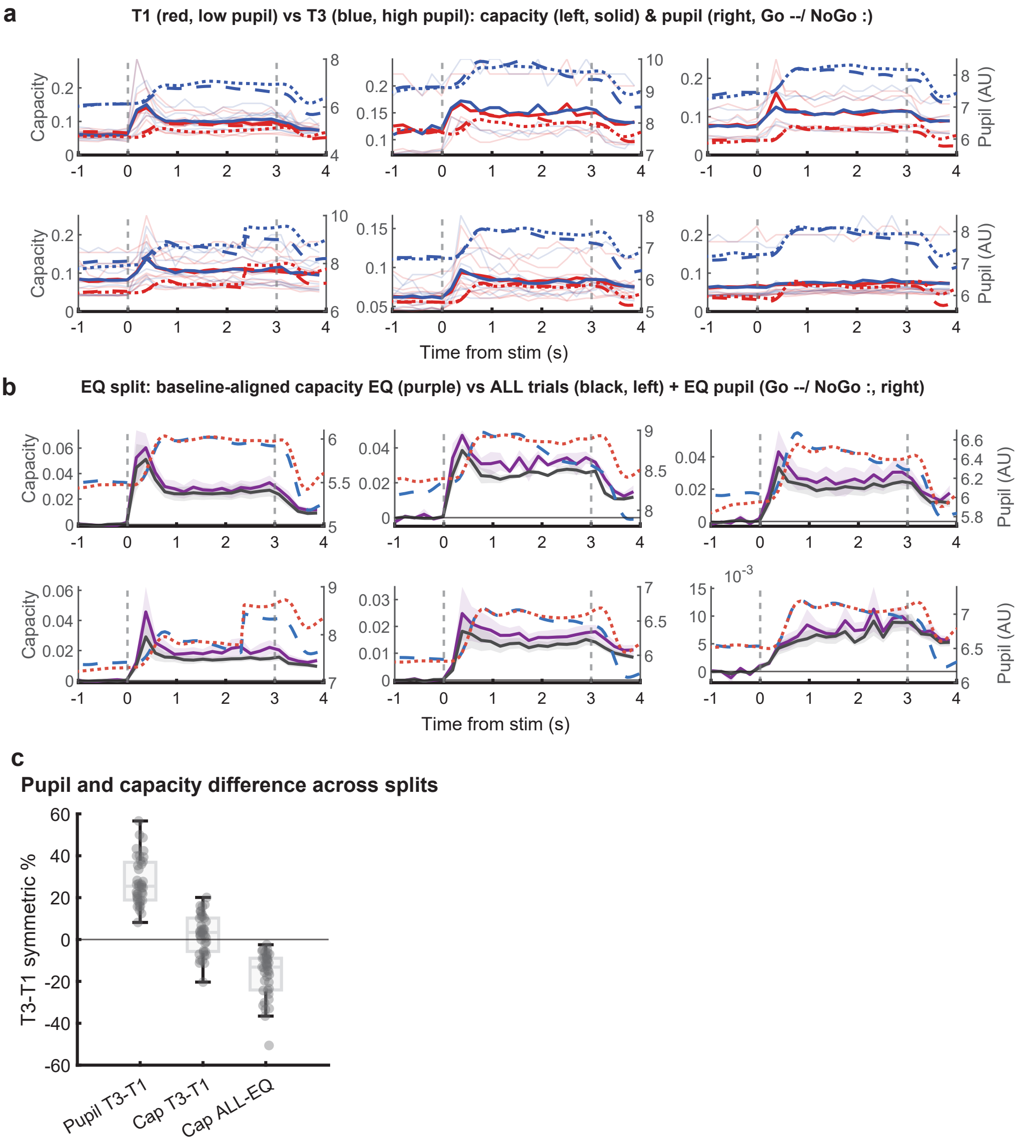


**Figure S9. Trial-evoked pupil size does not drive the capacity of the Trained representations. a.** Same analysis as fig. S8b, but splitting trials by their pupil size during the trial (1–2 s post-stimulus onset) instead of the pre-stimulus window. Left axis: capacity for the low (T1, red) and high (T3, blue) pupil tertiles, with thin lines showing individual sessions and thick lines the group averages. Right axis: average pupil traces of the Go (dashed) and NoGo (dotted) trials of each tertile. **b.** Same as fig. S8c for the 1–2 s split: capacity computed from the pupil-matched Go/NoGo trial subset (EQ, purple) compared with the capacity from all trials (black), both baseline-subtracted; shaded areas indicate s.e.m. Right axis: average pupil traces of the matched Go (dashed) and NoGo (dotted) trials. **c.** Same as fig. S8d: per-session percentage differences for pupil T3−T1, capacity T3−T1 and all-trials versus pupil-equalized capacity, all over the 1–2 s window. Boxes show median and interquartile range, whiskers the range, and dots individual sessions.

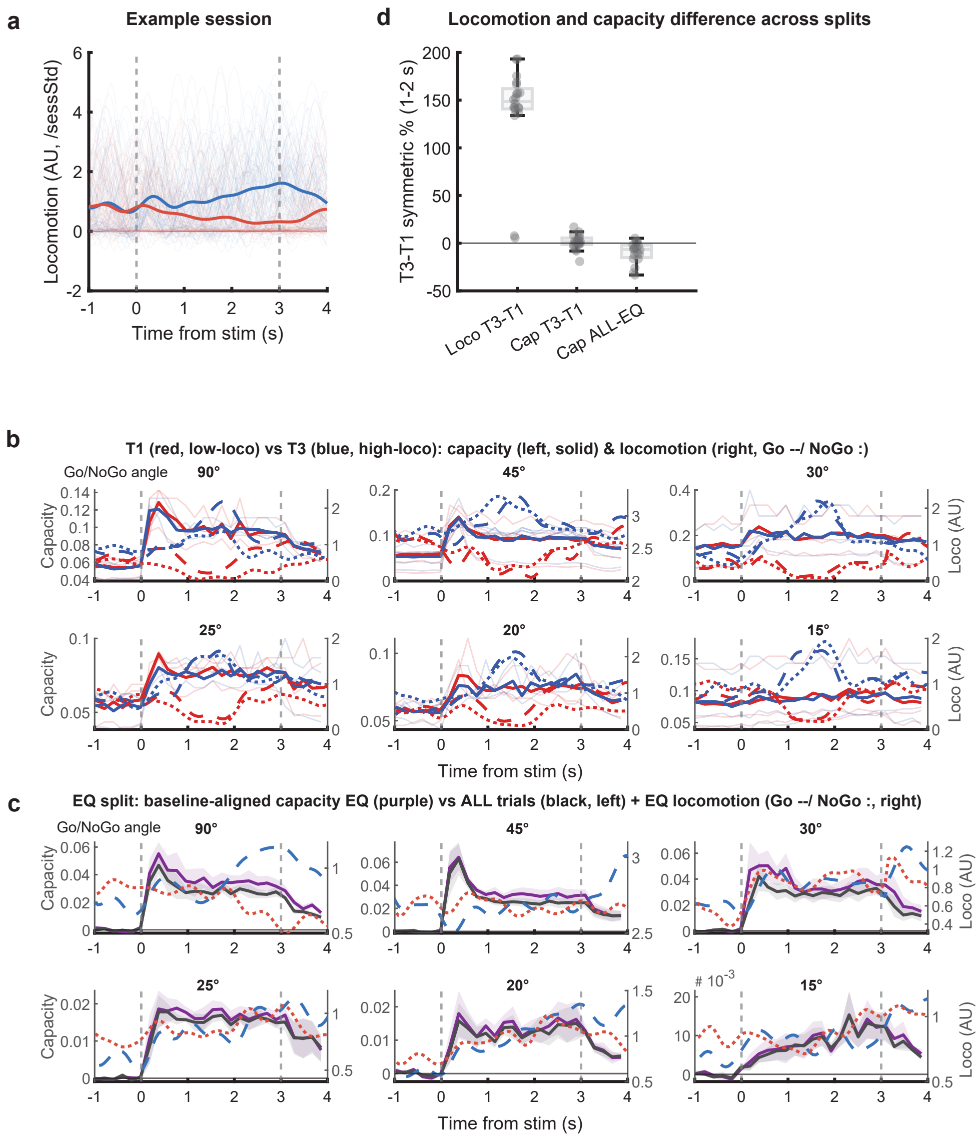


**Figure S10. Locomotion does not drive the capacity of the Trained representations. a.** Locomotion time course for an example session. Thin lines show single trials, thick lines the Go (blue) and NoGo (red) averages. Locomotion traces were low-pass filtered (1 Hz) and normalized by the session's standard deviation. Dashed lines indicate stimulus onset and offset. **b.** Same analysis as fig. S8b, but splitting trials by their locomotion during the trial (1–2 s post-stimulus onset). Left axis: capacity for the low (T1, red) and high (T3, blue) locomotion tertiles, with thin lines showing individual sessions and thick lines the group averages. Right axis: average locomotion traces of the Go (dashed) and NoGo (dotted) trials of each tertile. **c.** Same as fig. S8c for locomotion: capacity computed from the locomotion-matched Go/NoGo trial subset (EQ, purple) compared with the capacity from all trials (black), both baseline-subtracted; shaded areas indicate s.e.m. Right axis: average locomotion traces of the matched Go (dashed) and NoGo (dotted) trials. **d.** Same as fig. S8d: per-session symmetric percentage differences for locomotion T3−T1, capacity T3−T1 and all-trials versus locomotion-equalized capacity, all over the 1–2 s window. Boxes show median and interquartile range, whiskers the range, and dots individual sessions.

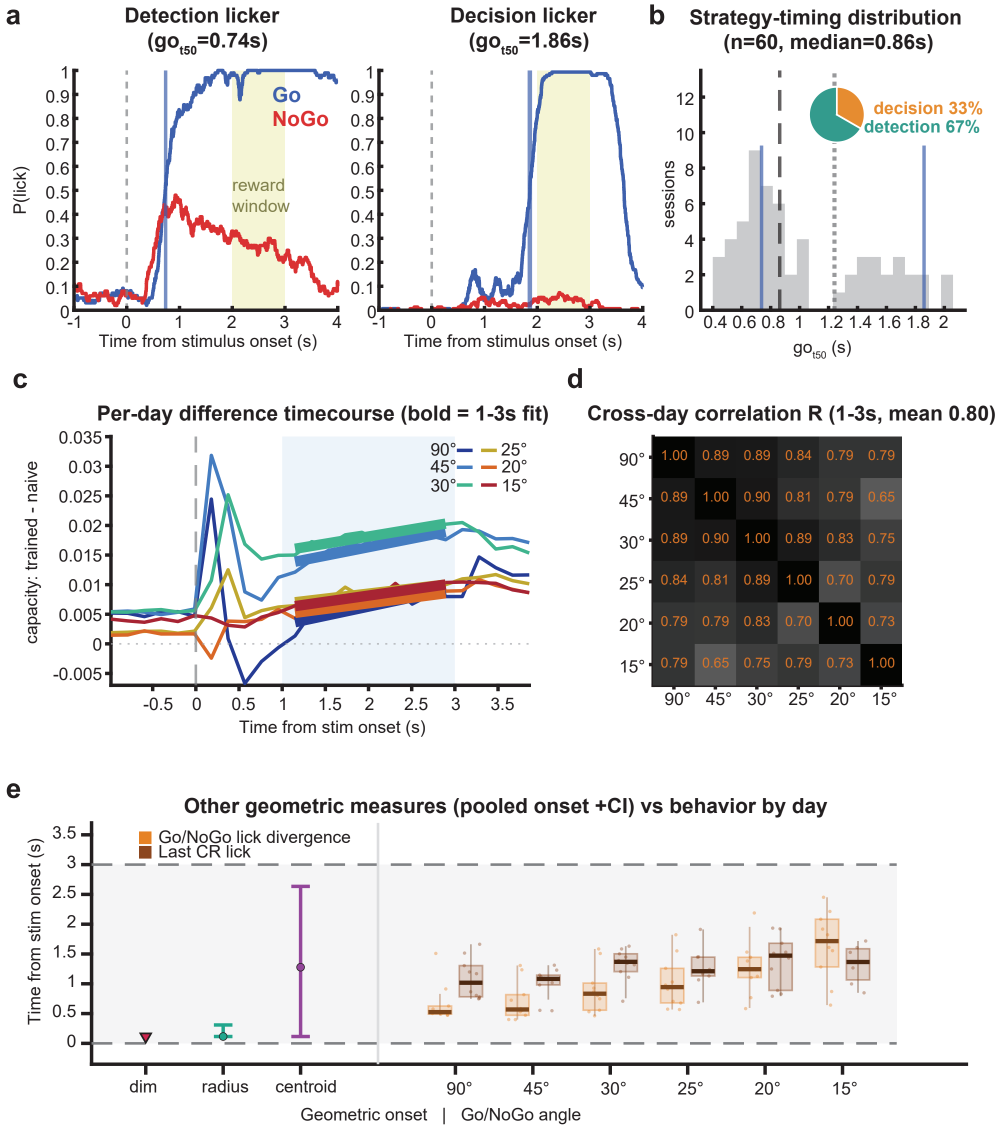


**Figure S11. Licking strategies, consistency of the capacity difference dynamics across Go/NoGo angles, and divergence timing of the geometric measures.** **a.** Lick probability time courses for Go (blue) and NoGo (red) trials in two example sessions illustrating the two licking strategies of the cohort: a "detection licker" (left) starting to lick shortly after stimulus onset regardless of its identity, and a "decision licker" (right) withholding licking until closer to the reward window (shaded). The licking onset is quantified as got50, the time to reach half of the peak Go lick probability (vertical blue line). The dashed line indicates stimulus onset. **b.** Distribution of got50 over all Trained sessions (n = 60, median 0.86 s). Vertical blue lines mark the two example sessions from a; the dashed line marks the cohort median and the dotted line the strategy split threshold. The pie chart indicates the proportion of sessions in each strategy (detection 67%, decision 33%). **c.** Time course of the Trained−Naive capacity difference computed separately for every Go/NoGo angle (colors). Bold lines show the linear fits over the 1–3 s window (shaded). **d.** Correlation matrix of the per-angle difference time courses from c over the 1–3 s window (mean R = 0.80), showing that the capacity difference follows similar dynamics at every Go/NoGo angle, allowing the sessions to be pooled for the estimation of the divergence onset. **e.** Same as fig. 6b but for the divergence onsets of the other geometric measures: naive-versus-trained onset of the dimensionality, radius and centroid distance differences (dots, pooled onset; bars, 95% bootstrap interval), compared with the per-Go/NoGo-angle distributions of the Go/NoGo lick divergence (orange) and last correct-rejection lick (brown) across Trained sessions. Boxes show median and interquartile range, whiskers the range, and dots individual sessions. Dashed lines indicate stimulus onset and offset.

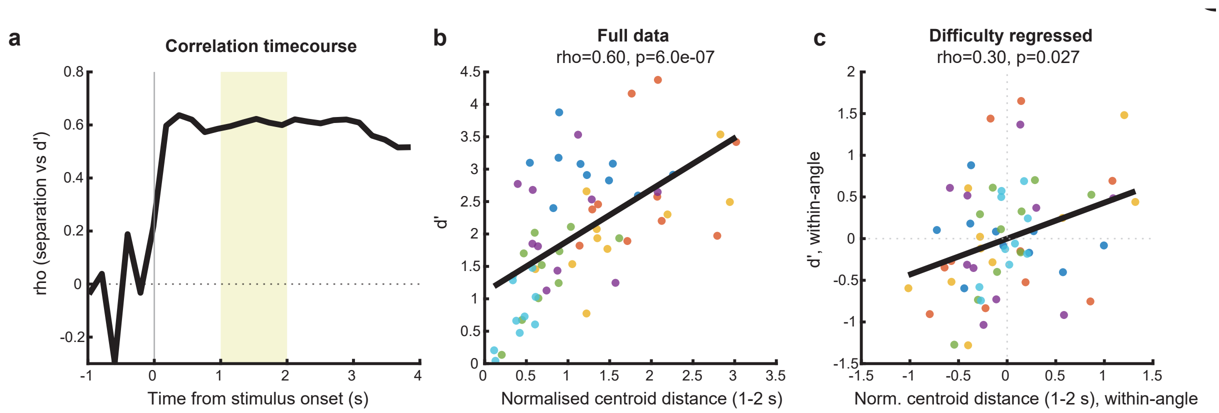


**Figure S12. Go/NoGo separation in the embedding space retains a within-difficulty correlation with performance. Same as fig. 6c–e but for the spread-normalized Go/NoGo separation in the PaCMAP embedding, computed as the distance between the Go and NoGo centroids divided by the mean of the two class radii, removing the arbitrary scale of each session's embedding. a.** Time course of the Spearman correlation between the normalized separation and behavioral sensitivity D′ across Trained sessions; the shaded band marks the 1–2 s window. **b.** D′ against the normalized separation averaged over 1–2 s, one dot per session, colored by Go/NoGo angle; the line is the linear fit. Spearman rho = 0.60, $p = 6.0\times10^{-7}$, n = 60. **c.** Same as b after regressing out task difficulty (within-recording-day-centered separation and D′). Partial Spearman rho = 0.30, p = 0.027.

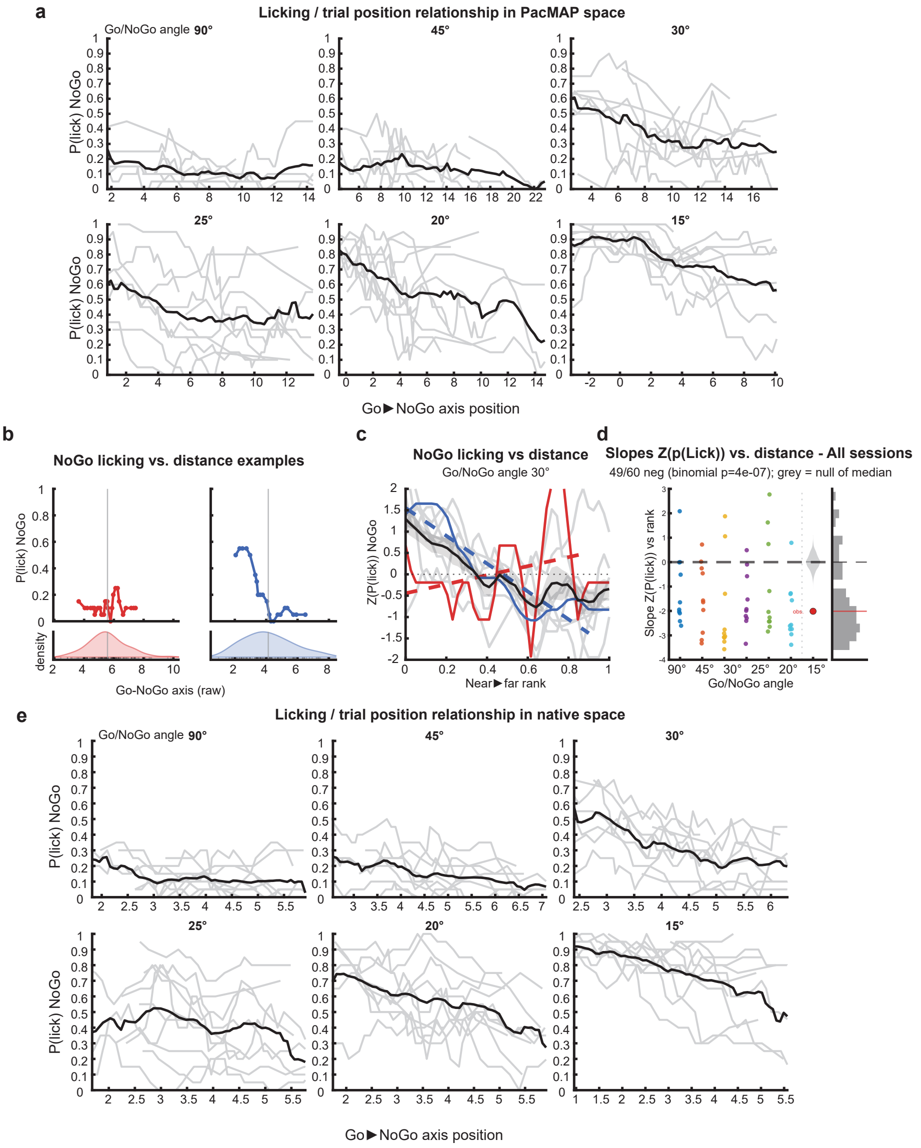


**Figure S13. Licking versus trial position along the Go to NoGo axis: raw positions and native space. a.** NoGo lick probability as a function of the raw (non rank-normalized) position along the Go to NoGo centroid axis in the PaCMAP embedding, for every Go/NoGo angle. Binned P(lick) on NoGo trials (sliding window of 20 trials, step 5); thin grey lines show individual sessions, black lines the across-session average. **b.** Same as fig. 6g but in the native space (no dimensionality reduction): NoGo lick probability as a function of the raw Go to NoGo axis position for two example sessions. Top: binned P(lick) on NoGo trials. Bottom: density of NoGo trials along the axis; ticks mark individual trials; the vertical line indicates the Go centroid. **c.** Same as fig. 6h but in the native space: NoGo licking versus axis position aggregated across Trained sessions at a Go/NoGo angle of 30°. Each session's binned NoGo lick probability was z-scored and plotted against the within-session near to far rank; thin grey lines show individual sessions, the black line the mean, and the red and blue lines the two example sessions from b, with dashed lines indicating their linear fits. **d.** Same as fig. 6i but in the native space: distribution of per-session slopes of z(P(lick)) versus near to far rank, by day and pooled. 49/60 slopes negative (binomial $p = 4\times10^{-7}$); median slope −2.01. The observed median (red line) is compared with the shuffled-null distribution (grey violin; permutation $p < 5\times10^{-4}$). Right: histogram of all session slopes. **e.** Same as a but in the native space.

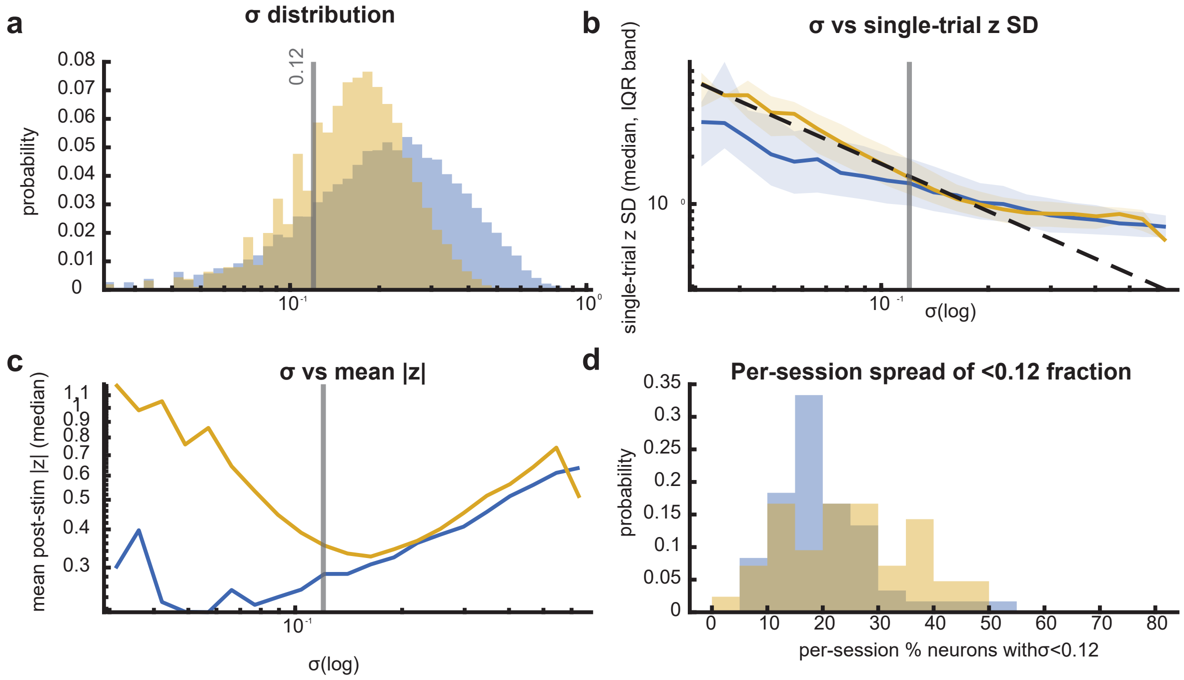


**Figure S14. Choice of the baseline standard deviation floor ($\sigma$ = 0.12). Diagnostics of the per-neuron baseline standard deviation ($\sigma$, integrated-APrE units) used for the baseline z-scoring, computed over the whole dataset (34,973 neurons, 102 sessions), in Trained (blue) and Naive (gold) animals. The vertical grey line indicates the 0.12 floor in all panels. a.** Distribution of the per-neuron baseline $\sigma$ (log scale). 20% (Trained) and 24% (Naive) of neurons fall below the floor. **b.** Single-trial z-score standard deviation (i.e., the trial-to-trial noise) as a function of $\sigma$ (median and IQR band). The dashed line shows the $1/\sigma$ relationship expected when the z-score spread is dominated by the estimation error on $\sigma$. Below the floor, the spread rises along this line, closely in Naive animals, in a more attenuated form in Trained animals, whose evoked activity is sparser. Above the floor, both groups flatten to the signal level. The floor was set at this knee (
textasciitilde24 baseline spikes per neuron,
textasciitilde10% relative error on the $\sigma$ estimate). **c.** Mean post-stimulus absolute z-score (i.e., the evoked signal) as a function of baseline $\sigma$. The low-$\sigma$ inflation is mainly expressed in Naive animals, whose low $\sigma$ baseline neurons had some activity during the stimulus. This mirrors panel b: for these same neurons, the trial-to-trial noise inflates with the same $1/\sigma$ profile as the signal, so the amplification rescales the two together. The floor therefore removes a group-asymmetric scale artifact (without excluding any neuron from the analyses), and prevents noisy large z-scores from contaminating distances. **d.** Distribution across sessions of the percentage of neurons with $\sigma$ below the floor.
